# Taxonomic distribution of SbmA/BacA and BacA-like antimicrobial peptide transporters suggests independent recruitment and convergent evolution in host-microbe interactions

**DOI:** 10.1101/2024.02.25.581009

**Authors:** Nicholas T. Smith, Amira Boukherissa, Kiera Antaya, Graeme W. Howe, Ricardo C Rodríguez de la Vega, Jacqui A. Shykoff, Benoît Alunni, George C. diCenzo

**Affiliations:** Department of Biology, Queen’s University, Kingston, ON, K7L 3N6, Canada; Department of Chemistry, Queen’s University, Kingston, ON, K7L 3N6, Canada; Institute for Integrative Biology of the Cell, CNRS, CEA, Université Paris-Saclay, 91198, Gif-sur-Yvette, France; Écologie Systématique et Évolution, CNRS, Université Paris-Saclay, AgroParisTech, 91198, Gif-sur-Yvette, France; Université Paris-Saclay, INRAE, AgroParisTech, Institut Jean-Pierre Bourgin (IJPB), 78000, Versailles, France

**Keywords:** SbmA, antimicrobial peptides, host-microbe interaction, peptide transport, convergent evolution, rhizobium-legume symbioses, pathogenesis

## Abstract

Small, antimicrobial peptides are often produced by eukaryotes to control bacterial populations in both pathogenic and mutualistic symbioses. These include proline-rich mammalian immune peptides and cysteine-rich peptides produced by legume plants in symbiosis with rhizobia. The fitness of the bacterial partner is dependent upon their ability to persist in the presence of these antimicrobial peptides. In the case of *Escherichia coli* and *Mycobacterium tuberculosis* pathogens and nitrogen-fixing legume symbionts (rhizobia), the ability to survive exposure to these peptides depends on peptide transporters called SbmA (also known as BacA) or BclA (for BacA-like). However, how broadly these transporters are distributed amongst bacteria, and their evolutionary history, is poorly understood. Here, we used hidden Markov models, phylogenetic analysis, and sequence similarity networks to examine the distribution of SbmA/BacA and BclA proteins across a representative set of 1,255 species from across the domain *Bacteria*. We identified a total of 71 and 177 SbmA/BacA and BclA proteins, respectively. Phylogenetic and sequence similarity analyses suggest that these protein families likely did not evolve from a common ancestor and that their functional similarity is instead a result of convergent evolution. *In vitro* sensitivity assays using the legume peptide NCR247 and several of the newly-identified BclA proteins confirmed that transport of antimicrobial peptides is a common feature of this protein family. Analysis of the taxonomic distribution of these proteins showed that SbmA/BacA orthologs were encoded only by species in the phylum *Pseudomonadota* and that they were primarily identified in just two orders: *Hyphomicrobiales* (class *Alphaproteobacteria*) and *Enterobacterales* (class *Gammaproteobacteria*). BclA orthologs were somewhat more broadly distributed and were found in clusters across four phyla. These included several orders of the phyla *Pseudomonadota* and *Cyanobacteriota*, as well as the order *Mycobacteriales* (phylum *Actinomycetota*) and the class *Negativicutes* (phylum *Bacillota*). Notably, many of the clades enriched for species encoding BacA or BclA orthologs also include many species known to interact with eukaryotic hosts in mutualistic or pathogenic interactions. Collectively, these observations suggest that SbmA/BacA and BclA proteins have been repeatedly co-opted to facilitate both mutualistic and pathogenic associations with eukaryotic hosts by allowing bacteria to cope with host-encoded antimicrobial peptides.

## INTRODUCTION

Rhizobia are a polyphyletic group of bacteria from the classes *Alphaproteobacteria* and *Betaproteobacteria* that can enter endosymbiotic relationships with leguminous plants. During the symbiosis, rhizobia reside within cells of a specialized structure called a root nodule, where the bacteria convert atmospheric N_2_ gas into ammonium through a process known as symbiotic nitrogen fixation (SNF). The fixed nitrogen is provided to the plant host in exchange for photosynthetically fixed carbon, allowing plant growth in otherwise nitrogen-limited conditions (1). As a result, SNF is frequently leveraged in agriculture in place of nitrogen fertilizers, which are both expensive and a major source of agricultural greenhouse gas emissions (2, 3).

Within nodules of legumes of the inverted-repeat lacking clade (IRLC) and Dalbergioid clade, rhizobia undergo a process called terminal bacteroid differentiation (TBD) prior to fixing nitrogen (4, 5) This involves bacterial cell enlargement, genome endoreduplication, increased membrane permeability, and major changes in gene expression and cellular metabolism (6–8). TBD is essential for SNF in these legumes and is driven by a family of small, plant-encoded proteins known as nodule-specific cysteine-rich (NCR) peptides, which target the intracellular symbiotic bacteria to promote TBD (9, 10). IRLC and Dalbergioid legumes encode between <10 and >700 NCR peptides that contain four, six, or eight conserved cysteine residues (11, 12) but otherwise share little sequence conservation and have isoelectric points (pI) that vary from 3 (anionic) to 11 (cationic) (11, 13). Cationic NCR peptides with pI values above nine display antimicrobial activity, likely due to interaction with the negatively-charged surface of bacterial plasma membranes resulting in membrane depolarization (14, 15). During symbioses with rhizobia, NCR peptides are thought to additionally interact with rhizobial cytosolic proteins, such as cell cycle regulators and cell division proteins, to promote TBD (15, 16).

NCR peptide-triggered TBD involves bacteria-encoded ABC transporters known as BacA (a homolog of *Escherichia coli* SbmA) and BclA (BacA-like); all rhizobial symbionts of IRLC and Dalbergioid legumes encode either a BacA or BclA protein, but to our knowledge, not both. BacA and BclA are inner membrane peptide transporters, differing primarily by the presence of an ATPase domain in BclA that is absent in BacA (17). ATP hydrolysis by the ATPase domain is essential for the transport activity of BclA, whereas BacA-mediated transport is driven by the proton-motive force (18, 19). Despite this difference, both BacA and BclA can import NCR peptides, and their loss renders rhizobia hypersensitive to NCR peptide exposure *in vitro* (17). Loss-of-function mutation of *bacA* or *bclA* results in several other free-living phenotypes, including increased resistance to the antibiotics bleomycin and gentamicin and, in some species, increased sensitivity to detergents and altered lipopolysaccharide modifications (20–26). In addition, rhizobia carrying loss-of-function *bacA* or *bclA* mutations rapidly die upon release into the nodules of IRLC or Dalbergioid legumes in an NCR peptide-dependent fashion and thus fail to differentiate (17, 27). On the other hand, *bacA* and *bclA* are not required for symbiosis with legumes that produce no NCR peptides (21). It has been hypothesized that the requirement of BacA or BclA for SNF in legume plants that induce TBD may be two-fold: (i) to move NCR peptides away from the cell membrane, thereby protecting the bacteria from their antimicrobial activities, and (ii) to transport the NCR peptides to their intracellular targets to promote TBD (17, 28, 29). While BacA and BclA are the primary transporters of NCR peptides in all rhizobia for which this trait has been studied, recent results suggest that another broad-specificity ABC transporter, the *Sinorhizobium meliloti* YejABEF protein, may also contribute to the import of NCR peptides and bacterial survival when grown in the presence of NCR peptides (30).

Orthologs of BacA and BclA have also been found in multiple pathogenic bacteria, including *E. coli* (where it is known as SbmA) (31), *Brucella abortus* (32), and *Mycobacterium tuberculosis* (33). Similar to how BacA or BclA is required for rhizobial survival in nodules of legume plants inducing TBD, BacA and BclA orthologs are required by *B. abortus* and *M. tuberculosis* for chronic infection of their eukaryotic hosts (32–34), likely by providing resistance to host-encoded antimicrobial peptides as these transporters are required for peptide transport and resistance *in vitro* (18, 19). Despite the conserved role of BacA and BclA as peptide transporters essential for beneficial and pathogenic interactions, orthologs vary in their ability to functionally replace each other. Whereas the *bacA* genes of *B. abortus* and *S. meliloti* complement the symbiotic defect of a *S. meliloti bacA* null mutant (20, 35), little to no complementation of the symbiotic defects is observed when the *S. fredii*, *R. leguminosarum*, or *Mesorhizobium loti bacA* are expressed in an *S. meliloti bacA* null mutant (23, 28). Likewise, the *bclA* genes of *M. tuberculosis* and *Bradyrhizobium* spp. cannot restore nitrogen fixation when expressed in a *S. meliloti bacA* null mutant (33, 36, 37). However, despite the lack of complementation of symbiotic phenotypes, most *bacA* and *bclA* orthologs still complement the gentamicin and/or bleomycin phenotypes of *S. meliloti bacA* null mutants (20, 23, 28, 33, 35–37). These results suggest that BacA and BclA orthologs display slight variations in their peptide substrate range or rate of transport (13).

The observation that BacA and BclA orthologs are found in diverse bacterial lineages suggests that these proteins may be widespread housekeeping proteins subsequently co-opted for host-bacterial interactions (38). However, no systematic study of the distribution of BacA and BclA orthologs on the bacterial tree exists. In addition, the evolutionary relationship between the BacA and BclA families remains to be elucidated. Here, we report the distribution of BacA and BclA orthologs in 1,255 bacterial species from across the bacterial domain. We found BacA orthologs exclusively within the phylum *Pseudomonadales* (syn. *Proteobacteria*), while BclA orthologs were predominately limited to the phyla *Pseudomonadales*, *Cyanobacteriota* (syn. *Cyanobacteria*), *Actinomycetota* (syn. *Actinomycetes*), and *Bacillota* (syn. *Firmicutes*). Expression of a subset of the newly identified BclA proteins in *S. meliloti* Δ*bacA* mutants confirmed that transport of antimicrobial peptides is a common property of the BclA protein family. The taxonomic distribution of SbmA/BacA and BclA, together with phylogenetic analysis of these proteins, leads us to suggest that the functional similarities between SbmA/BacA and BclA are a result of convergent evolution, and that these protein families have been repeatedly co-opted to help microbes cope with antimicrobial peptide exposure during host-microbe interactions.

## RESULTS

### Identification and classification of SbmA/BacA and BclA orthologs across the bacterial domain

To study the evolution and distribution of SbmA/BacA and BclA proteins, we searched the proteomes of 1,255 bacterial species, each belonging to a distinct genus, for proteins showing similarity to the SbmA/BacA-like family of PFAM (PF05992) (see Materials and Methods). This process led to the identification of 366 putative SbmA/BacA-like family proteins from 258 species. We further classified each of these 366 proteins into one of five protein classes according to Guefrachi and colleagues (36): SbmA/BacA, BclA, *Mycobacterium* BacA (a BclA-like family of proteins first identified in *M. tuberculosis*), ExsE (a related protein family involved in long-chain fatty acid transport), and the so-called *Bradyrhizobium* homologous clade (a related protein family with an unknown function). Initially, this classification was based on the use of hidden Markov models (HMMs), which was subsequently refined based on phylogenetic reconstruction and a sequence similarity network (SSN) as described below.

Using HMMs for these five protein classes, the 366 SbmA/BacA-like family proteins were initially classified into 79 SbmA/BacA proteins, 169 BclA proteins, 50 *Mycobacterium* BacA proteins, 52 ExsE proteins, and 16 *Bradyrhizobium* homologous clade proteins (Figure 1A). A maximum-likelihood phylogenetic analysis led to the identification of three primary monophyletic groups (Figure 1A). Clade A comprised 48 proteins and included most ExsE and all *Bradyrhizobium* homologous clade proteins, which we treated as the outgroup. Clade B included 34 proteins that were annotated as a mix of BclA and ExsE based on the HMMs. Clade C was the largest clade consisting of 284 proteins, and included most of the putative BclA, SbmA/BacA, and *Mycobacterium* BacA proteins.

**Figure 1.**
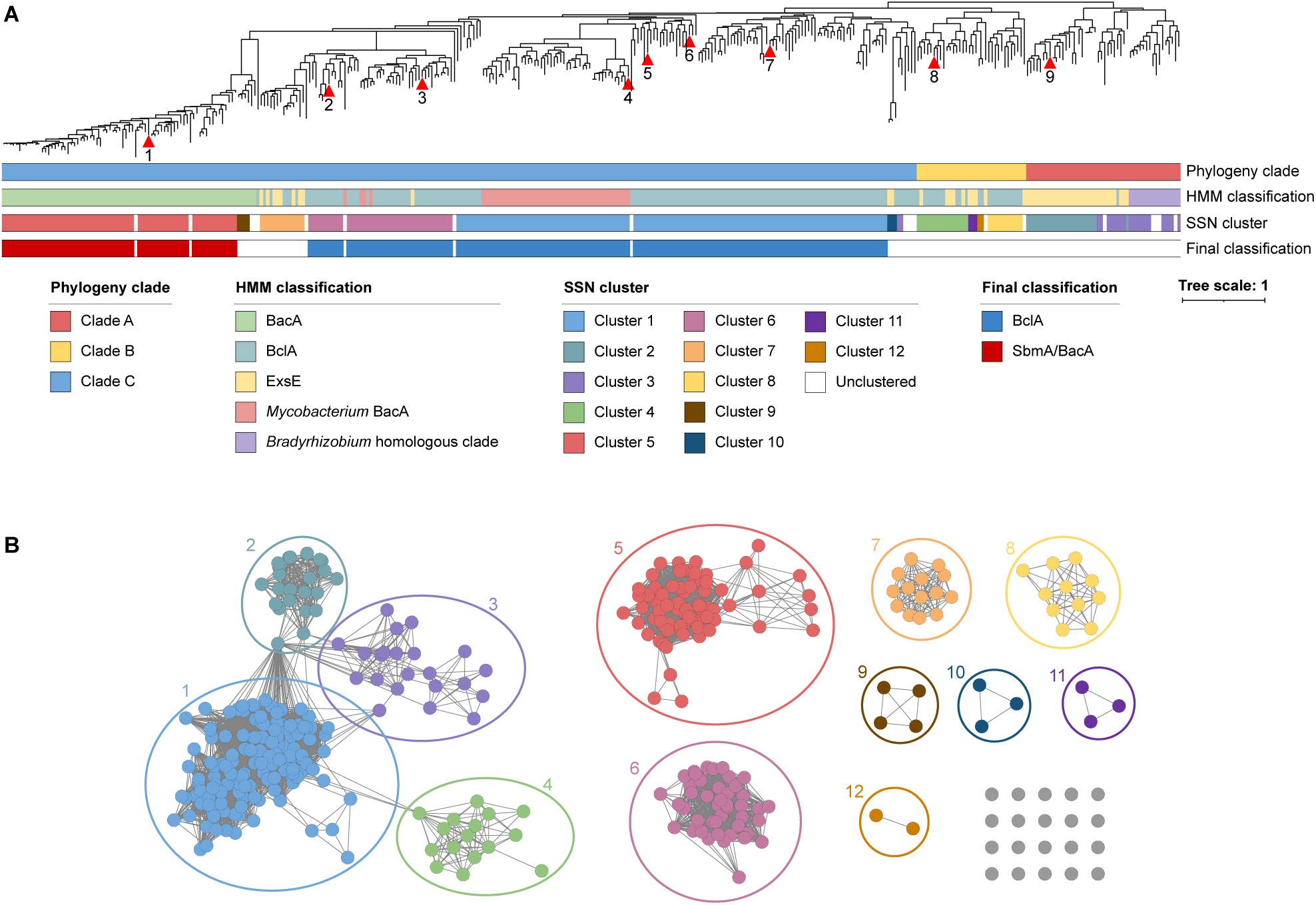
Sequence and phylogenetic analysis of SbmA/BacA-like proteins. **(A)** An unrooted maximum likelihood phylogeny of 366 SbmA/BacA-like proteins is shown. The scale bar represents the average number of amino acid substitutions per site. Red triangles indicate proteins whose corresponding genes were codon optimized and synthesized: 1 - *Polymorphum gilvum* BclA; 2 - *Synechococcus elongatus* BclA; 3 - *Cyanobacterium aponinum* BclA; 4 - *Basilea psittacipulmonis* BclA; 5 - *Succinivibrio dextrinosolvens* BclA; 6 - *Methylomusa anaerophila* BclA; 7 - *Polaromonas naphthalenivorans* BclA; 8 - *Eikenella exigua* BclA-like; 9 - *Phyllobacterium zundukense* ExsE. The bars beneath the phylogeny summarize the clustering and annotation of these proteins. The top bar indicates the phylogenetic clade to which each protein belongs. The second bar indicates the preliminary hidden Markov model (HMM) classification of each protein. The third bar indicates the cluster in the sequence similarity network that each protein belongs to. The bottom bar indicates which proteins were ultimately classified as SbmA/BacA (red) or BclA (blue). An interactive version of this phylogeny, with node support values, is provided through iTol (https://itol.embl.de/shared/1IAjjFrHYGLI9) while a Newick-formatted version of the phylogeny can be downloaded from GitHub (https://github.com/amira-boukh/SbmA_BacA_phylogenetic_distribution). **(B)** A sequence similarity network, calculated using EFI-EST, of 366 SbmA-BacA-like proteins is shown. Each node (the circles) represents one protein, while edges (the lines) represent sequence similarity between pairs of proteins above the threshold, with longer lines indicating lower similarity. Nodes are colour coded based on cluster.

Most of the putative BclA proteins from Clade C also form a single cluster in the SSN (Cluster 1; Figure 1B). We therefore conclude that the 133 proteins of Cluster 1 in the SSN represent true BclA orthologs. Notably, Cluster 1 of the SSN also includes 46 proteins annotated as *Mycobacterium* BacA, which also fall within Clade C in the phylogeny (Figure 1). This suggests that the *Mycobacterium* BacA proteins are not a distinct family from the BclA proteins, and that *Mycobacterium* BacA proteins should instead be referred to as BclA. On the other hand, a Clade C subclade of nine proteins with long branch lengths in the phylogeny is excluded from Cluster 1 of the SSN; instead, two of these proteins are found as part of Cluster 3 that predominantly consists of the *Bradyrhizobium* homologous clade proteins, three are found as a three-protein cluster (Cluster 10), and five are singletons. In addition, four of these nine proteins are from strains encoding a BclA protein belonging to Cluster 1. Taken together, we conclude that these nine proteins are not true BclA orthologs. Another subclade of Clade C consisting of 58 proteins is not part of Cluster 1 in the SSN but rather is largely found in two clusters (Clusters 6 and 7) of 44 and 14 proteins, respectively (Figure 1). Cluster 6 consists primarily of proteins from cyanobacteria, and 43 of the 44 proteins were classified as BclA or *Mycobacterium* BacA by the HMMs. In addition, the functional data described below suggests proteins of this cluster are functionally similar to known BclA proteins. We therefore conclude that proteins of Cluster 6 represent BclA orthologs. In contrast, eight of the 14 proteins of Cluster 7 were annotated as ExsE by the HMMs. The distinct clustering of Cluster 7 from Cluster 6, together with the HMM annotations, lead us to suggest that the proteins of Cluster 5 are unlikely to represent true BclA orthologs.

Consistent with the phylogenetic analysis, proteins of Clade B do not cluster with proteins of Clade C in the SSN (Figure 1B). Rather, the Clade B proteins are split across four clusters and two singletons. Nearly 1/3^rd^ (10 of 34) proteins of Clade B were annotated as ExsE by the initial HMM strategy, and many of the proteins of Clade B are from bacterial strains that also encode a putative SbmA/BacA or BclA of Clade C. Collectively, we interpret these results to indicate that Clade B proteins are not part of the BclA protein family, and that they instead represent a related but distinct protein family. This conclusion is also supported by the functional data presented below.

Lastly, all putative SbmA/BacA proteins formed a monophyletic group in the phylogeny (Figure 1A) and a monophyletic group of 71 of the 79 proteins form a single cluster (Cluster 5) in the SSN (Figure 1B). These results suggest that the 71 proteins of Cluster 5 and annotated as SbmA/BacA by the HMM strategy are likely true SbmA/BacA orthologs, and that all SbmA/BacA proteins evolved from a common ancestor. Although the SbmA/BacA proteins fell within Clade C in the phylogeny, the SbmA/BacA clade is connected to the rest of the tree via an unusually long branch, consistent with the distinct clustering of SbmA/BacA proteins in the SSN. The distinct clustering in the SSN, the long branch length, and the functional differences in transport (ATP-driven vs proton-driven) lead us to suggest that the SbmA/BacA and BclA protein families evolved independently and that their functional similarity is a result of convergent evolution.

In considering the different sources of information described above, we ultimately chose to select a final set of SbmA/BacA and BclA proteins based primarily on the SSN, resulting in the identification of 177 high-confidence BclA proteins (including the *Mycobacterium* BacA proteins) and 71 high-confidence BacA proteins (Figure 1A).

### *In vitro* functional analysis of diverse SbmA/BacA and BclA orthologs

To validate that the BclA and SbmA/BacA proteins identified through the *in silico* approach are functionally similar to known BclA and SbmA/BacA proteins, genes encoding nine of the identified proteins were synthesized. The proteins encoded by these genes included: one BacA protein, six BclA proteins including one previously classified as *Mycobacterium* BacA, one protein from Clade B (henceforth referred to as BacA-like), and one ExsE protein for comparison. The nine genes were then cloned into an expression vector and introduced into *S. meliloti* Δ*bacA* and *S. meliloti* Δ*bacA* Ω*yejA* mutants to test for complementation. Although the genes were codon-optimized for expression in *S. meliloti*, we cannot exclude the possibility that some proteins were not properly expressed or were not stably inserted into the *S. meliloti* inner membrane. Therefore, lack of complementation may reflect improper expression/localization of a protein rather than a lack of orthology. All strains showed similar growth in media lacking antimicrobial agents (Figure S1), indicating that differences in media supplemented with gentamicin (Gm) or NCR peptides reflect altered resistance phenotypes rather than general growth differences. In addition, we observed that the resistance phenotypes of the *S. meliloti* Δ*bacA* mutant complemented with the *S. meliloti bacA* gene *in trans* differed somewhat from wildtype *S. meliloti* (Figure S2), likely due to elevated expression of *bacA* in the complemented strain. Thus, for all *in vitro* phenotypic experiments, strains were compared to the *S. meliloti* Δ*bacA* mutant complemented with the *S. meliloti bacA* gene *in trans* rather than the wild type.

We first tested whether the nine genes could complement the Gm resistance phenotype of the *S. meliloti* Δ*bacA* mutant. As expected, the Δ*bacA* mutant was resistant to Gm, and reintroduction of the *S. meliloti bacA* gene *in trans* resulted in sensitivity to Gm (Figure 2A). Unexpectedly, introduction of the *Phyllobacterium zundukense exsE* gene resulted in intermediate complementation of the Gm resistance phenotype (Figure 2A), suggesting that transport of Gm is a broadly conserved function of the SbmA/BacA and related proteins, and is not specific to BclA or SbmA/BacA proteins. As a result, the impact of the nine genes on Gm resistance cannot be used to support the annotation of a protein specifically as BclA or SbmA/BacA; however, it is still a useful metric to test whether a SbmA/BacA-like protein is expressed and functional. Of the six *bclA* genes identified by our screen, three (from *Cyanobacterium aponinum*, *Synechococcus elongatus*, and *Succinivibrio dextrinosolvens*) complemented the Gm resistance phenotype at least as well as the known *bclA* gene of *Bradyrhizobium* sp. ORS285 (Figure 2A), confirming they are expressed and functional in *S. meliloti*. The other three *bclA* genes all displayed partial complementation to varying degrees (Figure 2B), suggesting they are expressed and functional but either have reduced ability to transport Gm or their expression or stability is sub-optimal. Likewise, the one BclA-like gene (from *Eikenella exigua*) displayed partial complementation of the Gm resistance phenotype (Figure 2B). On the other hand, introduction of the one *bacA* gene that we tested (from *Polymorphum gilvum*) completely failed to complement the Gm resistance phenotype of the *S. meliloti* Δ*bacA* mutant (Figure 2B), which we hypothesize is due to improper expression or stability of the protein rather than functional divergence.

**Figure 2.**
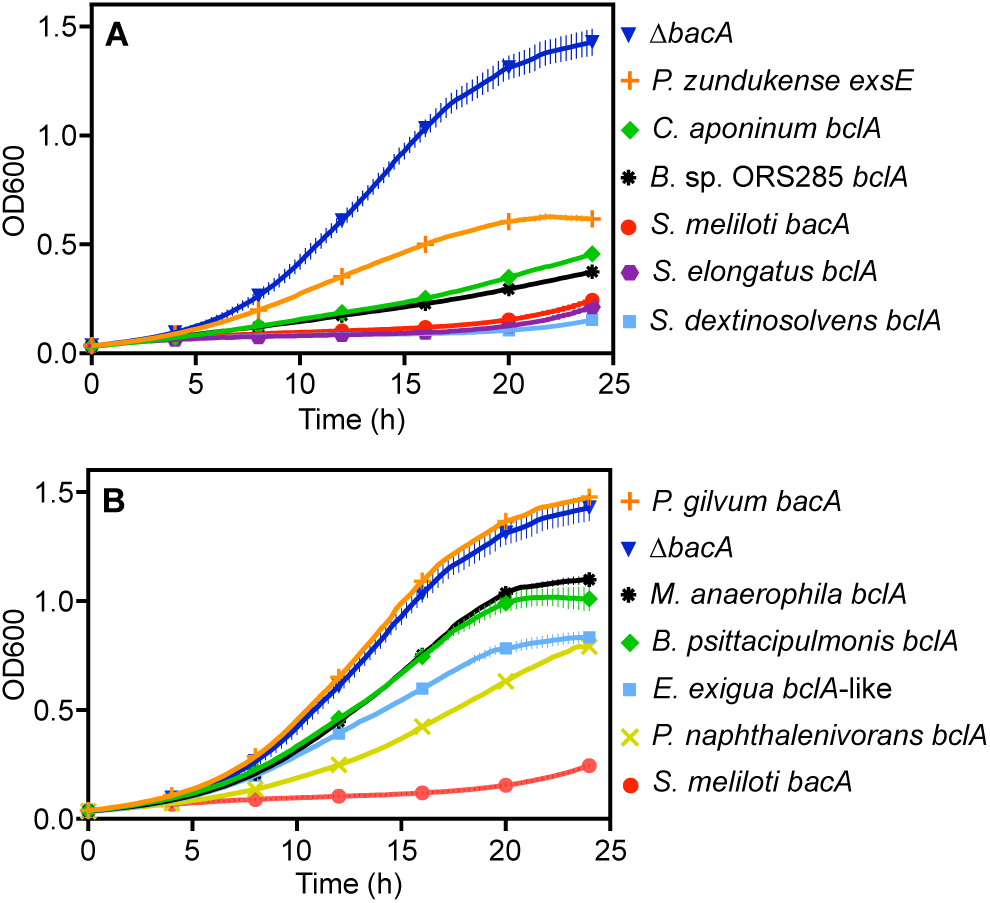
Gentamicin sensitivity assays. The growth of various *S. meliloti* strains, as measured by OD600, in the presence of 20 µg/mL of gentamicin is shown over a 24-hour period. Each point represents the mean of triplicate wells, with error bars depicting standard deviation. The Δ*bacA* strain represents the *S. meliloti* Δ*bacA* mutant carrying an empty vector, while all other strains are named according to the species of origin of the gene expressed *in trans* in the *S. meliloti* Δ*bacA* background. The experiment was replicated three independent times, and data from a representative experiment is shown. **(A)** Data is shown for genes exhibiting moderate to high level of complementation of the *S. meliloti* Δ*bacA* gentamicin resistance phenotype. **(B)** Data is shown for genes exhibiting low to moderate levels of complementation of the *S. meliloti* Δ*bacA* gentamicin resistance phenotype.

We next indirectly examined whether the nine proteins could transport eukaryotic antimicrobial peptides by measuring the impact of the proteins on the sensitivity of *S. meliloti* to the legume-encoded NCR peptide NCR247 (Figure 3); proteins transporting NCR247 are expected to show reduced sensitivity to this peptide. As expected, the *S. meliloti* Δ*bacA* single mutant and the Δ*bacA* Ω*yejA* double mutant were hypersensitive to NCR247 exposure, while introduction of the known *S. meliloti bacA* or *Bradyrhizobium* sp. ORS285 *bclA* genes *in trans* resulted in reduced sensitivity to NCR247 (Figure 3). Introduction of the *P. zundukense exsE* gene into the two mutants resulted in little to no complementation of the NCR247 hypersensitivity phenotypes (Figure 3), consistent with the transport of NCR peptides being specific to the SbmA/BacA and BclA family proteins and not a general property of these and related proteins. All three of the *bclA* genes showing strong complementation of the Gm resistance phenotype (two of which are from cyanobacteria) also showed good complementation of the NCR247 hypersensitivity phenotype (Figure 3), confirming the proteins encoded by these three genes are functionally similar to known BclA proteins. In addition, the *bclA* gene from *P. naphthalenivorans* strongly complemented the NCR247 hypersensitivity phenotypes of both strains despite only moderate complementation of the Gm resistance phenotype. Of the remaining two *bclA* genes, one (from *Methylomusa anaerophila*) displayed weak complementation of the NCR247 hypersensitivity (Figure 3) and varied in its level of complementation across trials (not shown), while one (from *Basilea psittacipulmonis*) failed to complement (Figure 3). Overall, the data for the six BclA proteins support that most BclA proteins are capable of transporting NCR peptides. On the other hand, the NCR247 sensitivity phenotypes of the strains expressing the BclA-like protein from *E. exigua* resembled the phenotypes of the strain expressing *P. zundukense exsE* (Figure 3), consistent with BclA-like proteins of Clade B (Figure 1) representing a different class of proteins from BclA. In accordance with the Gm resistance data, the *bacA* gene from *P. gilvum* largely failed to complement the NCR247 hypersensitivity phenotypes (Figure 3), potentially reflecting improper expression or stability of the encoded protein.

**Figure 3.**
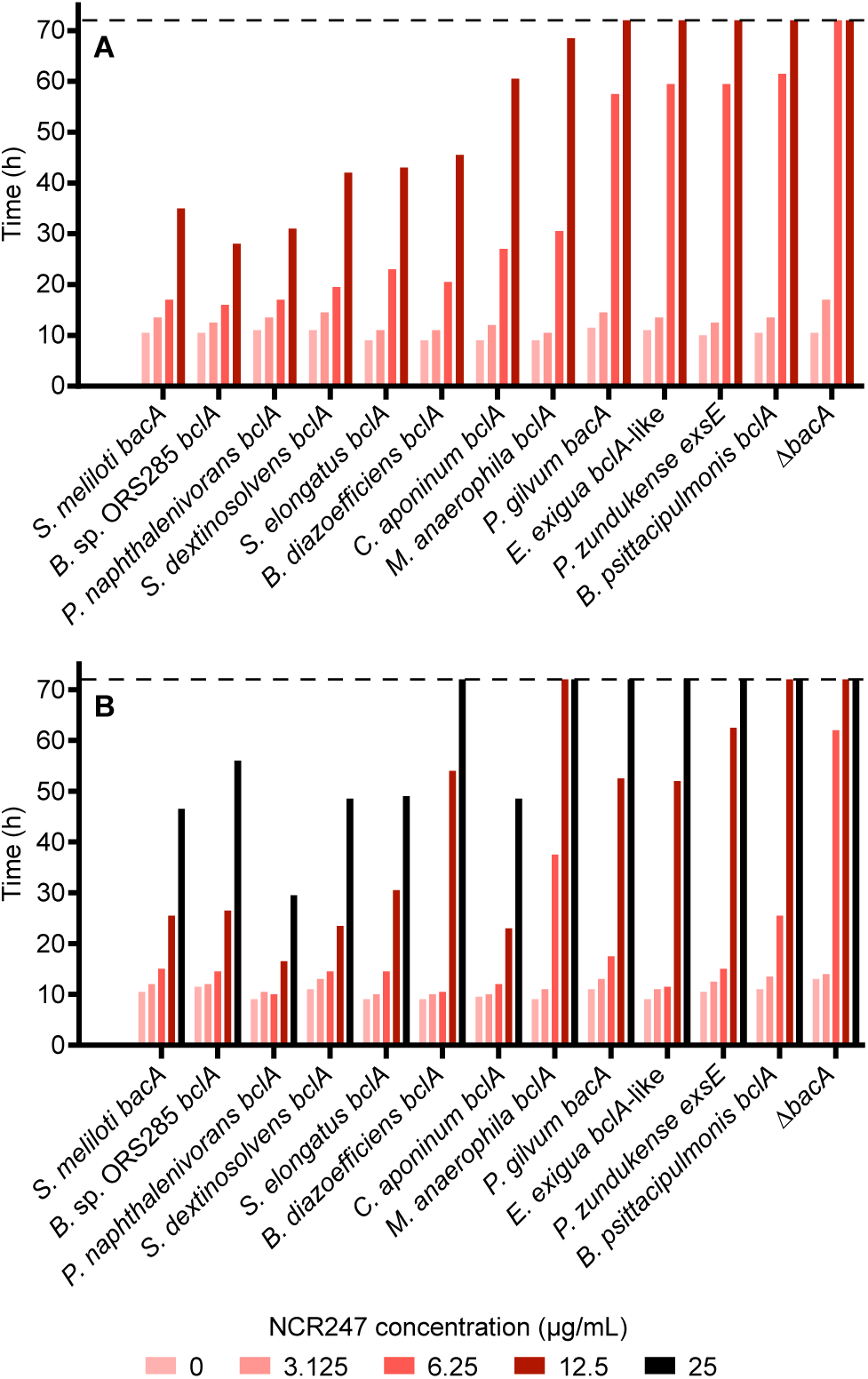
NCR247 sensitivity assays. The growth of various *S. meliloti* strains, as measured by OD600, in the presence of the antimicrobial peptide NCR247 is shown. Strains were grown in various concentrations of NCR247 as indicated by the shade of red or black. Bars represent the time required for the culture to reach an OD600 of 0.25. Values of 72 hours (indicated by the dashed line) indicate that the strain failed to reach an OD600 of 0.25 within the 72-hour growth period. The Δ*bacA* label represents the *S. meliloti* (A) Δ*bacA* or (B) Δ*bacA* Ω*yejA* mutant carrying an empty vector, while all other strains are named according to the species of origin of the gene expressed *in trans* in the *S. meliloti* (A) Δ*bacA* or (B) Δ*bacA* Ω*yejA* background. **(A)** Data is shown for the *S. meliloti* Δ*bacA* mutant and derivatives. **(B)** Data is shown for the *S. meliloti* Δ*bacA* Ω*yejA* mutant and derivatives.

### Analysis of the ability of BacA and BclA to support legume symbiosis

We additionally tested whether the nine proteins could complement the nitrogen-fixation defect of a *S. meliloti* Δ*bacA* mutant during symbiosis with *Medicago sativa* (alfalfa) or *Melilotus officinalis* (yellow-blossom sweet clover). As expected, the *S. meliloti* Δ*bacA* mutant formed small white nodules on both plants and failed to fix nitrogen, while re-introduction of the *S. meliloti bacA* gene *in trans* complemented the nitrogen-fixation phenotype (Table S1). All nine of the synthesized genes failed to complement the nitrogen-fixation phenotype (Table S1). As the same lack of complementation was observed for the known *bclA* gene of *Bradyrhizobium* sp. ORS285 (Table S1), these results suggest that most, if not all, BclA proteins are unable to support an effective symbiosis between *S. meliloti* and its host plants.

### Taxonomic distribution of SbmA/BacA and BclA orthologs across the domain ***Bacteria***

We next examined the taxonomic distribution of the 177 BclA and 71 SbmA/BacA proteins identified as described earlier. Remarkably, 100% and 78% of the identified SbmA/BacA and BclA proteins, respectively, are encoded by species of the phylum *Pseudomonadales* (syn. *Proteobacteria*) (Figure 4). As expected, most species encoding SbmA/BacA or BclA proteins encode only one or the other; only six of the 208 species encoding SbmA/BacA and/or BclA encode both, and in all six cases, both genes are carried by the chromosome.

**Figure 4.**
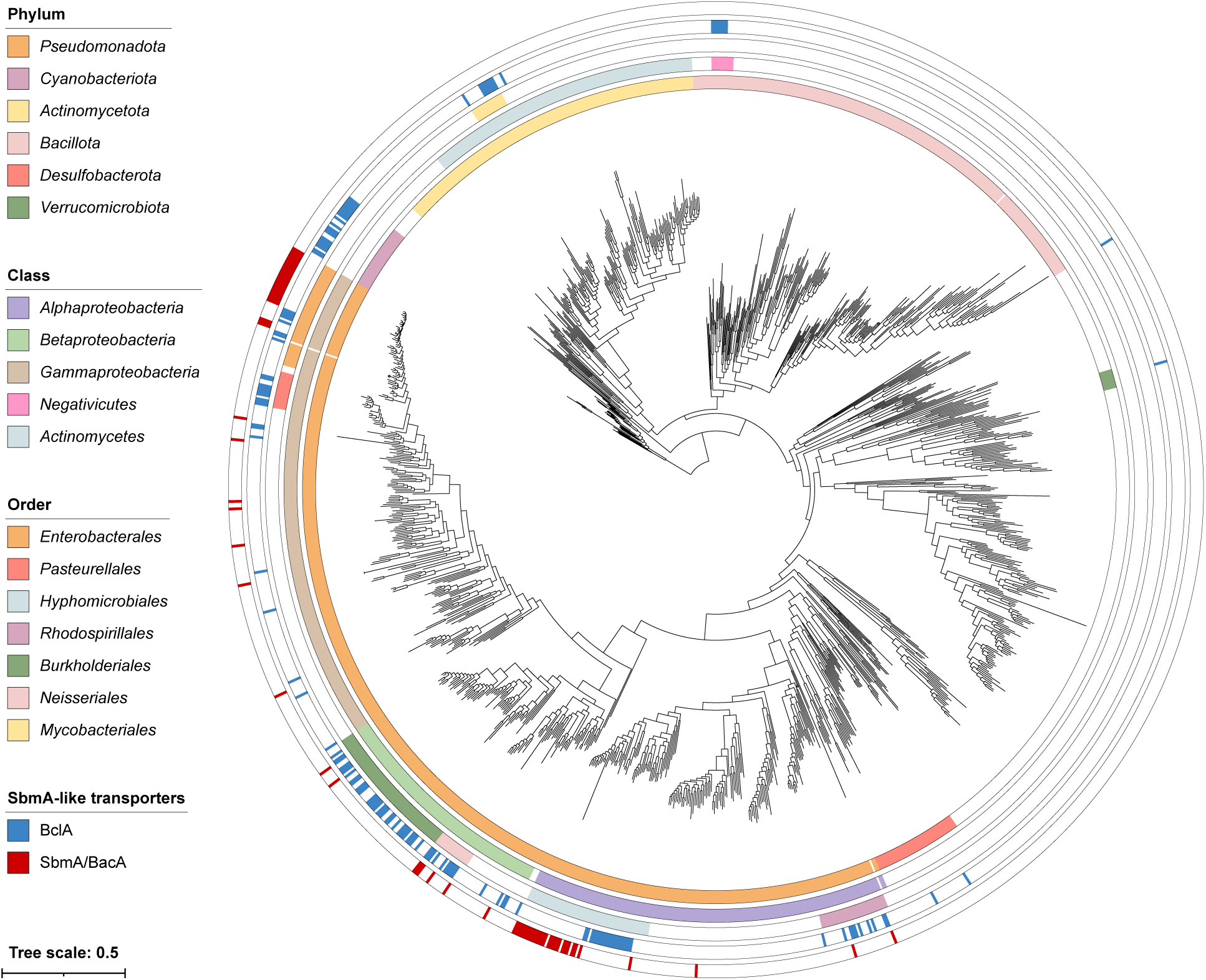
Taxonomic distribution of SbmA/BacA and BclA proteins in the domain *Bacteria*. An unrooted maximum likelihood phylogeny of 1,533 bacteria is shown, inferred from the concatenated protein alignments of 31 single-copy proteins. The scale bar represents the average number of amino acid substitutions per site. Three clades of intracellular symbionts/pathogens with long branch lengths were removed for presentation purposes; none of these taxa encode SbmA/BacA or BclA. The outer rings represent the following, starting from the inner ring: (i) the phylum that each strain belongs to, limited to phyla where at least one strain encodes SbmA/BacA or BclA; (ii) the class that each strain belongs to, limited to classes where at least one strain encodes SbmA/BacA or BclA and that are mentioned in the text; (iii) the class that each strain belongs to, limited to classes where at least one strain encodes SbmA/BacA or BclA and that are mentioned in the text; (iv) whether the strain encodes BclA (blue) or not (white); (v) whether the strain encodes SbmA/BacA (red) or not (white). An interactive version of this phylogeny, with node support values and without collapsing of any clades, is provided through iTol (https://itol.embl.de/shared/1IAjjFrHYGLI9) while a Newick-formatted version of the phylogeny can be downloaded from GitHub (https://github.com/amira-boukh/SbmA_BacA_phylogenetic_distribution).

Approximately 70% of the SbmA/BacA proteins are encoded by just two monophyletic groups of organisms, suggesting that SbmA/BacA was acquired at the base of each clade and then vertically transmitted. These two clades are a 24 species clade in the order *Enterobacterales* (all of which encode BacA) and a 29 species clade in the order *Hyphomicrobiales* (25 of which encode SbmA/BacA) (Figure 4). Interestingly, the SbmA/BacA proteins of the order *Enterobacterales* form a monophyletic group in the SbmA/BclA protein phylogeny (Figure S3). On the other hand, the minimal monophyletic clade encompassing all *Hyphomicrobiales* SbmA/BacA proteins also includes the *Enterobacterales* SbmA/BacA proteins (Figure S3). These results suggest that SbmA/BacA proteins of the order *Enterobacterales* were acquired through horizontal transfer from the order *Hyphomicrobiales*. The remaining 22 SbmA/BacA proteins not found within those two clades are distributed across the phylum *Psedomonadales* with no other major clustering observed. Overall, these results suggest that although SbmA/BacA proteins are widespread amongst subclades of the orders *Enterobacterales* (class *Gammaproteobacteria*) and *Hyphomicrobiales* (class *Alphaproteobacteria*), the taxonomic distribution of this protein family is otherwise limited.

BclA proteins show a somewhat broader taxonomic distribution than the SbmA/BacA proteins, although their distribution remains restricted to only a few phyla (Figure 4). Like SbmA/BacA, BclA was common in a subclade of the order *Hyphomicrobiales*, in which 19 of 21 species encoded BclA. Most of the other *Alphaproteobacteria* species encoding BclA belong to the order *Rhodospirillales*, in which nine of the 30 species encoded BclA. Within the *Gammaproteobacteria*, the taxon most enriched for BclA proteins was the order *Pasteurellales*, in which 10 of the 16 species encoded BclA. BclA was also abundant in the class *Betaproteobacteria*, unlike SbmA/BacA, and was particularly enriched in the orders *Burkholderiales* (32/60 species) and *Neisseriales* (10/17 species) compared to the orders *Nitrosomonadales* and *Rhodocyclales* (4/27 species across both orders). In contrast to SbmA/BacA, which was predicted to be encoded only by species of the phylum *Pseudomonadales*, there were three main clades of organisms predicted to encode BclA outside of the phylum *Pseudomonadales* (Figure 4). The largest of these were the phylum *Cyanobacteriota* (syn. *Cyanobacteria*), in which BclA was broadly distributed and found in 21 of the 31 species (∼67%). The other two main groups of organisms encoding BclA are a subclade of eight species (seven of which encode BclA) of the order *Mycobacteriales* (phylum *Actinomycetota* [syn. *Actinomycetes*]), and the class *Negativicutes* (phylum *Bacillota* [syn. *Firmicutes*]) in which seven of the ten species encode BclA orthologs (Figure 4).

## DISCUSSION

We identified 71 SbmA/BacA and 177 BclA orthologs from a search of the proteomes of 1,255 bacterial species. In total, 208 of the 1,255 species (16.6%) encoded at least one copy of SbmA/BacA and/or BclA, with only six of the 208 species (2.9%) encoding both SbmA/BacA and BclA. The observation that SbmA/BacA and BclA proteins were generally not encoded in the same proteome suggests that these protein families have similar biological roles. We also observed that the so-called “*Mycobacterium* BacA” proteins clustered with the BclA proteins in both the SSN and the protein phylogeny, leading us to conclude that the “*Mycobacterium* BacA” proteins are not distinct from BclA; we therefore reclassified the “*Mycobacterium* BacA” proteins as BclA for downstream analyses.

### Convergent evolution of the SmbA/BacA and BclA protein families

One of the objectives motivating this work was to gain insight into whether the SbmA/BacA and BclA protein families share common ancestry (e.g., that SbmA/BacA evolved from BclA, or vice versa) or whether they evolved independently and converged towards a similar function. The taxonomic distribution of SbmA/BacA and BclA proteins within the order *Hyphomicrobiales* is potentially suggestive of the former scenario. Excluding the deep-branching lineages, the order *Hyphomicrobiales* can be sub-divided into two sister clades; SbmA/BacA is widely distributed in one of these clades, while BclA is widely distributed in the other. This could suggest that the SbmA/BacA and BclA proteins of the order *Hyphomicrobiales* evolved from a common ancestral protein present in the ancestor of these clades. However, the *Hyphomicrobiales* SbmA/BacA and BclA proteins are polyphyletic in the BacA/BclA protein phylogeny, which instead suggests that the SbmA/BacA and BclA proteins of the order *Hyphomicrobiales* were independently acquired. The distinct clustering of the BclA and SbmA/BacA proteins in the SSN further supports independent evolutionary origins for these proteins, as does the notably long branch connecting the SbmA/BacA clade to the rest of the phylogeny. Moreover, we consider the differences in transport mechanisms of SbmA/BacA (proton gradient-driven) and BclA (ATP-driven) to be more easily explained if these protein families have separate evolutionary histories. Overall, we interpret the evidence as suggesting that the SbmA/BacA and BclA protein families evolved independently, and that their functional similarity is a result of convergent molecular evolution.

Twenty-eight of the BclA proteins were encoded by 21 cyanobacteria. These 28 proteins formed a distinct cluster in the SSN together with 15 non-cyanobacterial BclA proteins, raising the possibility that these proteins also evolved independently from the rest of the BclA proteins. While we cannot rule out this possibility, we consider the evidence to be insufficient to reach this conclusion at this time.

### The SbmA/BacA and BclA protein families are associated with eukaryotic host interaction

A second objective of this work was to determine how broadly SbmA/BacA and BclA proteins are distributed across the domain *Bacteria*. Contrary to our initial expectations, we found that both protein families display limited taxonomic distribution. SbmA/BacA orthologs were identified only in the phylum *Pseudomonadales*, with ∼89% of the identified BacA proteins being encoded by species of the classes *Alphaproteobacteria* and *Gammaproteobacteria*. A majority of the identified BclA proteins were also found in species of the phylum *Pseudomonadales* with a bias towards the *Betaproteobacteria*; however, BclA proteins were also common in the phylum *Cyanobacteriota*, the class *Negativicutes* (phylum *Bacillota*), and the order *Mycobacteriales* (phylum *Actinomycetota*). Interestingly, many of the clades enriched for species encoding SbmA/BacA or BclA orthologs also include many species known to interact with eukaryotic hosts in mutualistic or pathogenic interactions.

Forty-five of the 55 species (∼82%) of the alphaproteobacterial order *Hyphomicrobiales* encode SbmA/BacA and/or BclA; this increases to 45 of 50 species (90%) when excluding the deep-branching *Hyphomicrobiales* lineages. This order accounts for ∼79% of the alphaproteobacterial species encoding SbmA/BacA and/or BclA orthologs. Many members of the order *Hyphomicrobiales* are notable for their ability to interact with eukaryotic hosts. All alpha-rhizobia belong to the order *Hyphomicrobiales*, which also encompasses several plant and mammalian pathogens like *Agrobacterium* and *Brucella*, respectively (39). Similarly, ∼75% of the gammaproteobacterial BacA and BclA proteins are encoded by species in the orders *Enterobacterales* and *Pasteurellales*, in which 34 of 47 (∼72%; increasing to 81% when excluding a monophyletic group of five obligate endosymbionts) and 10 of 16 (∼62.5%) species encode BacA/BclA, respectively. The order *Enterobacterales* is well-known for including many plant (e.g., *Dickeya*, *Pantoea*) and animal/human (e.g., *Klebsiella*, *Yersinia*) pathogens (40). Likewise, the order *Pasteurellales* encompasses several animal/human pathogens (e.g., *Haemophilus*, *Pasteurella*) (41). In the class *Betaproteobacteria*, BclA and SbmA/BacA were significantly more common in the orders *Burkholderiales* and *Neisseriales* compared to the orders *Nitrosomonadales* and *Rhodocyclales*. The order *Burkholderiales* encompass all known beta-rhizobia as well as insect gut symbionts (e.g.., *Caballeronia*) and plant (e.g., *Ralstonia*) and animal/human (e.g., *Burkholderia*) pathogens (42, 43). The order *Neisseriales* encompasses many mammalian commensals but also some human pathogens (e.g., *Neisseria*) (44).

The phylum *Cyanobacteria* is the largest clade of organisms encoding BclA proteins outside of the phylum *Pseudomonadales*. To our knowledge, cyanobacteria are not pathogenic. However, many can form beneficial associations with diverse hosts, such as the nitrogen-fixing symbiosis between *Nostoc* and plants (45), the mutualistic relationship with fungi (forming lichens), and with sponges (46). The order *Mycobacteriales* includes important human and plant pathogens (e.g.,*Mycobacterium*, *Rhodococcoides*) (47), and opportunistic pathogens (e.g., *Mycolicibacterium*) (48). The class *Negativicutes* is poorly studied despite its peculiar nature, as these *Firmicutes* possess an outer membrane and a LPS (49). Nevertheless, this class is a common component of eukaryotic microbiomes and can cause human disease, including meningitis (50).

The observation that most taxonomic clades enriched for species encoding SbmA/BacA or BclA also contain many mutualistic and/or pathogenic organisms may suggest that eukaryotic host interaction is a driver of SbmA/BacA and BclA maintenance in these lineages. However, the data also suggests that these protein families may pre-date these species interactions. Assuming that SbmA/BacA was acquired by the common ancestor of the BacA-containing subclade of the order *Hyphomicrobiales*, the SbmA/BacA protein family potentially evolved in this lineage over 500 million years ago (51), which predates the evolution of legumes that are estimated to have evolved around 60 million years ago (52). Thus, SbmA/BacA could not have evolved in this lineage as a response to legume symbiosis. Accordingly, rhizobia that do not face NCR peptides in their legume host have a SbmA/BacA protein that can transport peptides (24) while not being able to complement the symbiotic defect of a *S. meliloti bacA* mutant (28). Rather, we hypothesize that SbmA/BacA originally evolved to fulfil another role and was subsequently co-opted to support legume symbiosis in rhizobia. Likewise, we hypothesize that BclA already existed in the *Bradyrhizobium* lineage prior to the evolution of legume symbiosis, and that this protein was independently co-opted for legume symbiosis in these organisms, mimicking the convergent evolution of NCR peptides in the IRLC and Dalbergioid legume families (12). Another role for SbmA/BacA proteins that may predate the evolution of symbiosis is inter and intraspecific competition, as highlighted by a study of phazolicin-producing rhizobia (53). Phazolicin is a narrow-spectrum antibiotic peptide that is produced by some rhizobial strains and that can kill other rhizobia after being imported by BacA and YejABEF transporters (53). More broadly, we hypothesize that BacA and BclA proteins did not evolve specifically for host interaction but rather were repeatedly co-opted to help bacteria survive exposure to host encoded antimicrobial peptides (e.g., NCR peptides in rhizobium-legume symbioses, and mammalian immune antimicrobial peptides during infection).

### Functional characterization of SbmA/BacA and BclA protein families

The abilities of several newly identified BclA proteins to complement the phenotypes of a *S. meliloti* Δ*bacA* mutant were tested to validate that these proteins were correctly annotated. *S. meliloti bacA* null mutants display increased gentamicin resistance compared to the wild type (20). Eight of the nine synthesized genes at least partially complemented the gentamicin resistance phenotype of a *S. meliloti* Δ*bacA* mutant, suggesting these eight proteins were expressed and at least partially functional in *S. meliloti*. Interestingly, even the gene encoding an ExsE ortholog partially complemented the gentamicin resistance phenotype, indicating that gentamicin transport is not specific to SbmA/BacA and BclA proteins but is a general property of these and related protein families. Gentamicin sensitivity assays are commonly used to characterize the function of rhizobial *bacA* orthologs and rhizobial *bacA* mutant alleles generated through site-directed mutagenesis (26). Although these assays are useful to identify null phenotypes, our results show that they do not probe a function unique to SbmA/BacA or BclA proteins and thus have limited value as a proxy to peptide transport or host interaction assays.

In addition to showing increased resistance to gentamicin, *S. meliloti* Δ*bacA* mutants show increased sensitivity to NCR peptides (17, 26). As the antimicrobial activity of NCR peptides is a result of their interaction with the cell envelope, it is thought that SbmA/BacA and BclA proteins provide resistance to NCR peptides by moving the peptides away from the cell envelope and into the cell (17, 28). SbmA/BacA and BclA proteins have also been shown to transport other antimicrobial peptides, including mammalian antimicrobial peptides such as Bac7 (19, 33, 36, 54). As expected, only the proteins annotated as BclA were capable of effectively complementing the sensitivity of *S. meliloti* Δ*bacA* and *S. meliloti* Δ*bacA* Ω*yejA* mutants to the NCR peptide NCR247. Of the six newly-identified BclA proteins that were tested, four repeatedly demonstrated good levels of complementation; these proteins were from *P. naphthalenivorans* (class *Betaproteobacteria*), *S. dextrinosolvens* (class *Gammaproteobacteria*), *S. elongatus* (phylum *Cyanobacteriota*), and *C. aponinum* (phylum *Cyanobacteriota*). The other two, from *M. anaerophila* (class *Negativicutes*) and *B. psittacipulmonis* (class *Betaproteobacteria*), showed weak and variable or little to no complementation, respectively. However, there are thousands of distinct NCR peptides encoded across the legume family (11), and thus the inability of a transporter to transport NCR247 does not mean that it is unable to transport other NCR peptides, or mammalian antimicrobial peptides. Indeed, *S. meliloti yejA* mutants show increased sensitivity to the peptide NCR280 but not NCR247 (30). Regardless, these results support that the ability to transport antimicrobial peptides, including NCR peptides, is a general property of bacterial SbmA-like proteins.

### Conclusions

In summary, we identified 208 bacterial species encoding SbmA/BacA or BclA. These species were not equally distributed across the domain *Bacteria*; instead, SbmA/BacA proteins were found only in the phylum *Pseudomonadota*, while BclA proteins were primarily found within a subset of families across four phyla. Our analyses suggest that the SbmA/BacA and BclA protein families arose independently and that their functional similarity is a result of convergent evolution rather than shared ancestry. Our data also support the hypothesis that SbmA/BacA and BclA proteins have been repeatedly co-opted to facilitate both mutualistic and pathogenic associations with eukaryotic hosts by allowing bacteria to cope with host-encoded antimicrobial peptides.

## MATERIALS AND METHODS

### Bacterial strains and growth conditions

The bacterial strains used in this study are listed in Table S2. *E. coli* strains were cultured at 37 °C using Lysogeny Broth (LB; 10 g/L tryptone, 5 g/L yeast extract, 5 g/L NaCl). *S. meliloti* strains were grown at 28 °C using either LBmc (LB supplemented with 2.5 mM CaCl_2_ and 2.5 mM MgSO_4_), YEB (0.5% beef extract, 0.1% yeast extract, 0.5% peptone, 0.5% sucrose, 0.04% MgSO_4_ 7H_2_O, pH 7.5), or MM9 minimal medium (2% MOPS-KOH, 1.92% NH_4_Cl, 0.35% NaCl, 0.2% KH_2_PO_4_, 0.2% MgSO_4_, 0.05% CaCl_2_, 0.05% Biotin, 0.0004% CoCl_2_, 0.38% FeCl_3_, 1% Glucose, 1% Na_2_-succinate). Antibiotics were added as appropriate and included: ampicillin (Amp; 100 µg/mL), kanamycin (Km; 100 µg/mL), streptomycin (Sm; 200 or 500 µg/mL), spectinomycin (Sp; 50 µg/mL), and tetracycline (Tc; 5 µg/mL). Antibiotic concentrations were generally halved for liquid cultures.

### Cloning of ***bacA***, ***bclA***, and ***exsE*** homologs

Ten vectors encoding putative *bacA*, *bclA*, or *exsE* genes, codon optimized for *S. meliloti* 1021 and flanked by XbaI and BamHI recognition sites, were produced by Twist Biosciences (Table S2, Dataset S1). Each gene was PCR amplified from the plasmids using Q5 polymerase (New England Biolabs; NEB) with the primers 5’-GAAGTGCCATTCCGCCTGACC and 5’-CACTGAGCCTCCACCTAGCC. The resulting amplicons were individually digested with XbaI/BamHI and ligated into XbaI/BamHI-digested expression vector pRF771 (55). Plasmids were sequence verified via Illumina sequencing (151 bp paired-end reads) at SeqCenter (Pittsburg, PA, USA), after which reads were aligned to the expected template sequences using bowtie2 version 2.5.0 (56) and alignments visualized using the Integrative Genomics Viewer version 2.12.3 (57).

### Transfer of plasmids to ***S. meliloti***

All plasmids of interest were transferred to a *S. meliloti* Δ*bacA* mutant via triparental matings using the helper strains *E. coli* MT616 or *E. coli* HB101, as described previously (37, 58). Transconjugants were recovered through plating of mating spots on LBmc Sm^200^ Tc or YEB Sm^500^ Tc plates. Likewise, plasmids were transferred to a *S. meliloti* Δ*bacA* Ω*yejA* double mutant via triparental mating as described previously (37), with transconjugants recovered on YEB Sm^500^ Tc Km Sp plates. All transconjugants were streak purified three times prior to use.

### Gentamicin sensitivity assays

Gentamicin sensitivity assays were performed largely as described previously (28). Briefly, overnight cultures of *S. meliloti*, grown in LBmc Sm^100^ Tc, were washed and resuspended in LBmc to an optical density at 600 nm (OD600) of 1.0. Ten µL aliquots of the cell suspensions were added to triplicate wells of a 96-well plate and mixed with 190 µL of LBmc with or without 20 µg/mL of gentamicin (Gm). A Gm concentration of 20 µg/mL was chosen for the assays based on preliminary sensitivity assays (Figure S4). Plates were tape-closed to prevent evaporation and then incubated at 30°C with maximal shaking in a BioTek Synergy H1 plate reader for 24 hours. OD600 measurements were collected every 15 minutes using the Gen5 software (Agilent Technologies).

### NCR247 sensitivity assays

NCR sensitivity assays were performed largely as described previously (30). Briefly, overnight cultures of *S. meliloti*, grown in MM9 minimal media, were washed and resuspended in MM9 to an OD600 of 1.0. Cell suspensions were then diluted to an OD600 of 0.05 and 145 µL transferred to the wells of 96-well plates and mixed with 5 µL of an NCR247 solution to reach final concentrations of 50, 25, 12.5, 6.25, 3.125, and 0 µg/mL of NCR247. Plates were incubated at 28°C with shaking (180 rpm) in a Tecan Spark plate reader for 72 hours, and OD_600_ measurements were taken every 30 minutes and processed using the SparkControl software (Tecan).

### Plant symbiotic assays

Seeds of *M. sativa* cv. Algonquin (alfalfa) and *M. officinalis* (yellow-blossom sweet clover) (Speare Seeds Limited; Harriston, Ontario, Canada) were surface-sterilized and germinated on water agar plates for two nights in the dark, as described previously (13). Leonard assemblies were prepared as described before (13), with a 1:1 (w/w) mixture of vermiculite and silica sand in the top pot, 250 mL Jensen’s medium (59) in the bottom pot, and a cotton wick connecting the pots, and then autoclaved. Five seedlings were sown per pot, and assemblies were incubated for two nights. Assemblies were next inoculated in triplicate with 1 x 10^8^ CFU of *S. meliloti* per assembly. Plants were grown in a Conviron growth chamber with an 18-hour photoperiod, 300 µmol/s of light, 21 °C daytime temperature, and 17 °C nighttime temperature. After 30 days, plant shoots were collected and dried at 60 °C for six nights prior to weighing.

### Phylogenetic analysis of BacA and BclA proteins

GenBank files corresponding to 3498 RefSeq bacterial genomes with ‘complete’ genome assemblies were downloaded from the National Center for Biotechnology Information (NCBI) Genome Database. A subset of the genomes was prepared by collecting genomes from one representative genome per genus, using the genome from the first species per genus when sorted alphabetically. The phylogenetic analyses were then repeated twice: once using all 3498 RefSeq bacterial genomes and once using the reduced set of 1255 genomes (Dataset S2). As the results were similar, we only present results generated using the reduced dataset.

BacA, BclA, and related proteins were extracted from the bacterial proteomes using a modified version of an existing in-house pipeline (60). The seed alignment of the SbmA/BacA-like family, consisting of eight sequences, was downloaded from PFAM (PF05992), and a hidden Markov model (HMM) built using the hmmbuild function of HMMER version 3.3 (61). Separately a HMM database was built by combining (i) the complete PFAM-A version 31.0 HMM database, (ii) the complete TIGERFAM version 15.0 HMM database, (iii) HMMs built from the seed alignments of PRK11098 (105 sequences in the seed alignment) and COG1133 (nine sequences in the seed alignment) downloaded from NCBI’s Conserved Domain Database, and (iv) HMMs built for each of the BacA (15 sequences), BclA (5 sequences), *Mycobacterium* BacA (10 sequences), ExsE (6 sequences), and *Bradyrhizobium* homologous clade (7 sequences) proteins used in the phylogenetic analysis of Guefrachi et al. 2015 (36). Next, the hmmsearch function of HMMER was used to search all bacterial proteomes using the PF05992 (SbmA/BacA-like family) HMM. All hmmsearch hits were then scanned against the full HMM database using the hmmscan function of HMMER. Each protein was annotated according to the top-scoring HMM from this search.

Proteins annotated as BacA, BclA, *Mycobacterium* BacA, ExsE, or *Bradyrhizobium* homologous clade were extracted and aligned using Clustal Omega version 1.2.4 (62), hmmalign from HMMER (61), and MAFFT version 7.453 (63), and alignment quality assessed with T-COFFEE version 13.45 (64). Poor quality regions of the best scoring alignment (Clustal Omega) were removed using trimAl version 1.4 with the automated1 option (65), and then used as input for maximum likelihood phylogeny inference using IQ-TREE2 version 2.2.0 (66) with the LG+F+I+I+R9 model. The LG+F+I+I+R9 model was used as it was identified as the best-scoring model by the IQ-TREE2 implementation of ModelFinder (67) based on Bayesian information criterion (BIC), with model search limited to the LG, WAG, JTT, Q.pfam, JTTDCMut, DCMut, VT, PMB, BLOSUM62, and Dayhoff models. Branch supports were assessed in IQ-TREE using Shimodaira-Hasegawa-like approximate likelihood ratio test (SH-aLRT) (68) and an ultrafast bootstrap analysis, with both metrics calculated from 1000 replicates. All phylogenies created in this study were visualized with the iTOL web server (69).

### Sequence similarity network analysis

A sequence similarity network (SSN) was constructed for the 366 proteins identified using the HMM approach described above. The SSN was constructed using the online Enzyme Function Initiative’s Enzyme Similarity Tool (EFI-EST; https://efi.igb.illinois.edu/efi-est/) (70, 71) with an alignment score threshold of 115, corresponding to an approximate sequence ID ≥ 35%. The resulting network was visualized using Cytoscape version 3.10.1 (72).

### Multilocus sequence analysis

A bacterial species phylogeny was produced for the 1,253 representative bacterial species using an adaptation of an existing in-house pipeline (60); two of the 1,255 downloaded genomes were excluded as they encoded none of the marker genes. First, orthologs of 31 highly-conserved, single-copy proteins (DnaG, Frr, InfC, NusA, Pgk, PyrG, RplA, RplB, RplC, RplD, RplE, RplF, RplK, RplL, RplM, RplN, RplP, RplS, RplT, RpmA, RpoB, RpsB, RpsC, RpsE, RpsI, RpsJ, RpsK, RpsM, RpsS, SmpB, Tsf) were identified in the 1,253 proteomes using the AMPHORA2 pipeline (73). Each group of orthologs was individually aligned using MAFFT (63) and trimmed using trimAl with the option (65). The protein alignments were then concatenated and used as input for ModelFinder as implemented in IQ-TREE2, and the best scoring model was identified based on BIC. IQ-TREE2 was then used to infer a maximum likelihood phylogeny from the concatenated alignment using the LG+I+I+R10 model. Branch supports were assessed in IQ-TREE using the Shimodaira-Hasegawa-like approximate likelihood ratio test (SH-aLRT) [22] and ultrafast jackknife analysis with a subsampling proportion of 40%, with both metrics calculated from 1000 replicates.

## Data availability

All genome sequences used in this work were previously published, and the assembly accessions are provided in Dataset S2. Likewise, all protein sequences included in Figure 1 are provided in Dataset S1. Newick formatted phylogenies used to create Figures 1 and 4 are available through GitHub (https://github.com/amira-boukh/SbmA_BacA_phylogenetic_distribution). All code to repeat the analyses in this study is also available through GitHub (https://github.com/amira-boukh/SbmA_BacA_phylogenetic_distribution).

## ACKNOWLEDGEMENTS

This work was supported by Natural Sciences and Engineering Research Council of Canada (NSERC) Discovery Grants to G.C.D. and G.W.H.. N.T.S. was supported, in part, by an R. S. McLaughlin Fellowship from Queen’s University and a Wicked Ideas grant from Queen’s University to G.W.H and G.C.D.. A.B. benefited from a Ph.D. contract in the frame of the CNRS 80|PRIME – 2021 program and was partially supported by a Mitacs Globalink Research Award. B.A. benefited from a French State grant (Saclay Plant Sciences, reference n° ANR-17-EUR-0007, EUR SPS-GSR) under a France 2030 program (reference n° ANR-11-IDEX-0003).

## Notes

https://github.com/amira-boukh/SbmA_BacA_phylogenetic_distribution

https://itol.embl.de/shared/1IAjjFrHYGLI9

### Competing Interest Statement

The authors have declared no competing interest.

## REFERENCES

1. Poole P, Ramachandran V, Terpolilli J. 2018. Rhizobia: From saprophytes to endosymbionts. Nature Reviews Microbiology 16:291–303.

2. Menegat S, Ledo A, Tirado R. 2022. Greenhouse gas emissions from global production and use of nitrogen synthetic fertilisers in agriculture. Sci Rep 12:14490.

3. Peoples MB, Brockwell J, Herridge DF, Rochester IJ, Alves BJR, Urquiaga S, Boddey RM, Dakora FD, Bhattarai S, Maskey SL, Sampet C, Rerkasem B, Khan DF, Hauggaard-Nielsen H, Jensen ES. 2009. The contributions of nitrogen-fixing crop legumes to the productivity of agricultural systems. Symbiosis 48:1–17.

4. Alunni B, Gourion B. 2016. Terminal bacteroid differentiation in the legume−rhizobium symbiosis: nodule-specific cysteine-rich peptides and beyond. New Phytologist 211:411–417.

5. Haag AF, Mergaert P. 2020. Terminal bacteroid differentiation in the *Medicago*–*Rhizobium* interaction – a tug of war between plant and bacteria, p. 600–616. In The Model Legume Medicago truncatula. John Wiley & Sons, Ltd.

6. Mergaert P, Uchiumi T, Alunni B, Evanno G, Cheron A, Catrice O, Mausset A-E, Barloy-Hubler F, Galibert F, Kondorosi A, Kondorosi E. 2006. Eukaryotic control on bacterial cell cycle and differentiation in the Rhizobium–legume symbiosis. Proceedings of the National Academy of Sciences 103:5230–5235.

7. Roux B, Rodde N, Jardinaud M-F, Timmers T, Sauviac L, Cottret L, Carrère S, Sallet E, Courcelle E, Moreau S, Debellé F, Capela D, de Carvalho-Niebel F, Gouzy J, Bruand C, Gamas P. 2014. An integrated analysis of plant and bacterial gene expression in symbiotic root nodules using laser-capture microdissection coupled to RNA sequencing. Plant J 77:817–837.

8. diCenzo GC, Tesi M, Pfau T, Mengoni A, Fondi M. 2020. Genome-scale metabolic reconstruction of the symbiosis between a leguminous plant and a nitrogen-fixing bacterium. 1. Nat Commun 11:2574.

9. Van De Velde W, Zehirov G, Szatmari A, Debreczeny M, Ishihara H, Kevei Z, Farkas A, Mikulass K, Nagy A, Tiricz H, Satiat-Jeunemaître B, Alunni B, Bourge M, Kucho KI, Abe M, Kereszt A, Maroti G, Uchiumi T, Kondorosi E, Mergaert P. 2010. Plant peptides govern terminal differentiation of bacteria in symbiosis. Science 327:1122–1126.

10. Kereszt A, Mergaert P, Montiel J, Endre G, Kondorosi É. 2018. Impact of Plant Peptides on Symbiotic Nodule Development and Functioning. Frontiers in Plant Science 9:1026.

11. Montiel J, Downie JA, Farkas A, Bihari P, Herczeg R, Bálint B, Mergaert P, Kereszt A, Kondorosi É. 2017. Morphotype of bacteroids in different legumes correlates with the number and type of symbiotic NCR peptides. Proceedings of the National Academy of Sciences 114:5041–5046.

12. Czernic P, Gully D, Cartieaux F, Moulin L, Guefrachi I, Patrel D, Pierre O, Fardoux J, Chaintreuil C, Nguyen P, Gressent F, Da Silva C, Poulain J, Wincker P, Rofidal V, Hem S, Barrière Q, Arrighi J-F, Mergaert P, Giraud E. 2015. Convergent Evolution of Endosymbiont Differentiation in Dalbergioid and Inverted Repeat-Lacking Clade Legumes Mediated by Nodule-Specific Cysteine-Rich Peptides. Plant Physiol 169:1254–1265.

13. Huang R, Snedden WA, diCenzo GC. 2022. Reference nodule transcriptomes for *Melilotus officinalis* and *Medicago sativa* cv. Algonquin. Plant Direct 6:e408.

14. Szerencsés B, Gácser A, Endre G, Domonkos I, Tiricz H, Vágvölgyi C, Szolomajer J, Howan DHO, Tóth GK, Pfeiffer I, Kondorosi É. 2021. Symbiotic NCR Peptide Fragments Affect the Viability, Morphology and Biofilm Formation of *Candida* Species. Int J Mol Sci 22:3666.

15. Tiricz H, Szücs A, Farkas A, Pap B, Lima RM, Maróti G, Kondorosi É, Kereszt A. 2013. Antimicrobial nodule-specific cysteine-rich peptides induce membrane depolarization-associated changes in the transcriptome of *Sinorhizobium meliloti*. Applied and Environmental Microbiology 79:6737– 6746.

16. Penterman J, Abo RP, De Nisco NJ, Arnold MFF, Longhi R, Zanda M, Walker GC. 2014. Host plant peptides elicit a transcriptional response to control the *Sinorhizobium meliloti* cell cycle during symbiosis. Proceedings of the National Academy of Sciences 111:3561–3566.

17. Haag AF, Baloban M, Sani M, Kerscher B, Pierre O, Farkas A, Longhi R, Boncompagni E, Hérouart D, Dall’Angelo S, Kondorosi E, Zanda M, Mergaert P, Ferguson GP. 2011. Protection of *Sinorhizobium* against host cysteine-rich antimicrobial peptides is critical for symbiosis. PLoS Biology 9:e1001169.

18. Ghilarov D, Inaba-Inoue S, Stepien P, Qu F, Michalczyk E, Pakosz Z, Nomura N, Ogasawara S, Walker GC, Rebuffat S, Iwata S, Heddle JG, Beis K. 2021. Molecular mechanism of SbmA, a promiscuous transporter exploited by antimicrobial peptides. Science Advances 7:eabj5363.

19. Runti G, Lopez Ruiz M del C, Stoilova T, Hussain R, Jennions M, Choudhury HG, Benincasa M, Gennaro R, Beis K, Scocchi M. 2013. Functional characterization of SbmA, a bacterial inner membrane transporter required for importing the antimicrobial peptide Bac7. J Bacteriol 195:5343–5351.

20. Ichige A, Walker GC. 1997. Genetic analysis of the *Rhizobium meliloti bacA* gene: Functional interchangeability with the *Escherichia coli sbmA* gene and phenotypes of mutants. J Bacteriol 179:209–216.

21. Karunakaran R, Haag AF, East AK, Ramachandran VK, Prell J, James EK, Scocchi M, Ferguson GP, Poole PS. 2010. BacA Is Essential for Bacteroid Development in Nodules of Galegoid, but not Phaseoloid, Legumes. J Bacteriol 192:2920–2928.

22. Tan X-J, Cheng Y, Li Y-X, Li Y-G, Zhou J-C. 2009. BacA is indispensable for successful *Mesorhizobium*-*Astragalus* symbiosis. Appl Microbiol Biotechnol 84:519–526.

23. Maruya J, Saeki K. 2010. The *bacA* Gene Homolog, mlr7400, in *Mesorhizobium loti* MAFF303099 is Dispensable for Symbiosis with *Lotus japonicus* but Partially Capable of Supporting the Symbiotic Function of *bacA* in *Sinorhizobium meliloti*. Plant and Cell Physiology 51:1443–1452.

24. Ardissone S, Kobayashi H, Kambara K, Rummel C, Noel KD, Walker GC, Broughton WJ, Deakin WJ. 2011. Role of BacA in lipopolysaccharide synthesis, peptide transport, and nodulation by *Rhizobium* sp. strain NGR234. J Bacteriol 193:2218–2228.

25. Ferguson GP, Datta A, Carlson RW, Walker GC. 2005. Importance of unusually modified lipid A in *Sinorhizobium* stress resistance and legume symbiosis. Mol Microbiol 56:68–80.

26. LeVier K, Walker GC. 2001. Genetic analysis of the *Sinorhizobium meliloti* BacA protein: Differential effects of mutations on phenotypes. J Bacteriol 183:6444–6453.

27. Glazebrook J, Ichige A, Walker GC. 1993. A *Rhizobium meliloti* homolog of the *Escherichia coli* peptide-antibiotic transport protein SbmA is essential for bacteroid development. Genes and Development 7:1485–1497.

28. DiCenzo GC, Zamani M, Ludwig HN, Finan TM. 2017. Heterologous complementation reveals a specialized activity for BacA in the *Medicago*-*Sinorhizobium* meliloti symbiosis. Molecular Plant-Microbe Interactions 30:312–324.

29. Farkas A, Maróti G, Dürgo H, Györgypál Z, Lima RM, Medzihradszky KF, Kereszt A, Mergaert P, Kondorosi É. 2014. *Medicago truncatula* symbiotic peptide NCR247 contributes to bacteroid differentiation through multiple mechanisms. Proceedings of the National Academy of Sciences. 111:5183–5188.

30. Nicoud Q, Barrière Q, Busset N, Dendene S, Travin D, Bourge M, Le Bars R, Boulogne C, Lecroël M, Jenei S, Kereszt A, Kondorosi E, Biondi EG, Timchenko T, Alunni B, Mergaert P. 2021. *Sinorhizobium meliloti* Functions Required for Resistance to Antimicrobial NCR Peptides and Bacteroid Differentiation. mBio 12:10.1128/mbio.00895-21.

31. Laviña M, Pugsley AP, Moreno F. 1986. Identification, mapping, cloning and characterization of a gene (*sbmA*) required for microcin B17 action on *Escherichia coli* K12. J Gen Microbiol 132:1685– 1693.

32. LeVier K, Phillips RW, Grippe VK, Roop RM, Walker GC. 2000. Similar requirements of a plant symbiont and a mammalian pathogen for prolonged intracellular survival. Science 287:2492– 2493.

33. Domenech P, Kobayashi H, Levier K, Walker GC, Iii CEB. 2009. BacA, an ABC Transporter Involved in Maintenance of Chronic Murine Infections with *Mycobacterium tuberculosis*. J Bacteriol 191:477–485.

34. Li G, Laturnus C, Ewers C, Wieler LH. 2005. Identification of Genes Required for Avian *Escherichia coli* Septicemia by Signature-Tagged Mutagenesis. Infection and Immunity 73:2818.

35. Wehmeier S, Arnold MFF, Marlow VL, Aouida M, Myka KK, Fletcher V, Benincasa M, Scocchi M, Ramotar D, Ferguson GP. 2010. Internalization of a thiazole-modified peptide in *Sinorhizobium meliloti* occurs by BacA-dependent and -independent mechanisms. Microbiology 156:2702–2713.

36. Guefrachi I, Pierre O, Timchenko T, Alunni B, Barrière Q, Czernic P, Villaécija-Aguilar JA, Verly C, Bourge M, Fardoux J, Mars M, Kondorosi E, Giraud E, Mergaert P. 2015. *Bradyrhizobium* BclA Is a Peptide Transporter Required for Bacterial Differentiation in Symbiosis with *Aeschynomene* Legumes. Molecular Plant-Microbe Interactions 28:1155–1166.

37. Barrière Q, Guefrachi I, Gully D, Lamouche F, Pierre O, Fardoux J, Chaintreuil C, Alunni B, Timchenko T, Giraud E, Mergaert P. 2017. Integrated roles of BclA and DD-carboxypeptidase 1 in *Bradyrhizobium* differentiation within NCR-producing and NCR-lacking root nodules. Sci Rep 7:9063.

38. Arnold MFF, Haag AF, Capewell S, Boshoff HI, James EK, Donald RM, Mair I, Mitchell AM, Kerscher B, Mitchell TJ, Mergaert P, Barry IE, Scocchi M, Zanda M, Campopiano DJ, Ferguson GP. 2013. Partial complementation of *Sinorhizobium meliloti baca* mutant phenotypes by the *Mycobacterium tuberculosis* BacA protein. J Bacteriol 195:389–398.

39. diCenzo GC, Yang Y, Young JPW, Kuzmanović N. 2023. Refining the taxonomy of the order *Hyphomicrobiales* (*Rhizobiales*) based on whole genome comparisons of over 130 genus type strains. bioRxiv 10.1101/2023.11.15.567303.

40. de la Maza LM, Pezzlo MT, Bittencourt CE, Peterson EM. 2020. Introduction to *Enterobacterales*, p. 91–102. *In* Color Atlas of Medical Bacteriology. John Wiley & Sons, Ltd.

41. Garrity GM, Bell JA, Lilburn T. 2005. *Pasteurellales* ord. nov., p. 850–912. *In* Brenner, DJ, Krieg, NR, Staley, JT, Garrity, GM, Boone, DR, De Vos, P, Goodfellow, M, Rainey, FA, Schleifer, K-H (eds.), Bergey’s Manual® of Systematic Bacteriology: Volume Two The Proteobacteria Part B The Gammaproteobacteria. Springer US, Boston, MA.

42. Voronina OL, Kunda MS, Ryzhova NN, Aksenova EI, Semenov AN, Lasareva AV, Amelina EL, Chuchalin AG, Lunin VG, Gintsburg AL. 2015. The Variability of the Order Burkholderiales Representatives in the Healthcare Units. Biomed Res Int 2015:680210.

43. Dobritsa AP, Samadpour M. 2016. Transfer of eleven species of the genus Burkholderia to the genus Paraburkholderia and proposal of *Caballeronia* gen. nov. to accommodate twelve species of the genera Burkholderia and Paraburkholderia. International Journal of Systematic and Evolutionary Microbiology 66:2836–2846.

44. Chen S, Rudra B, Gupta RS. 2021. Phylogenomics and molecular signatures support division of the order *Neisseriales* into emended families *Neisseriaceae* and *Chromobacteriaceae* and three new families *Aquaspirillaceae* fam. nov., *Chitinibacteraceae* fam. nov., and *Leeiaceae* fam. nov. Systematic and Applied Microbiology 44:126251.

45. Bergman B, Johansson C, Söderbäck E. 1992. The *Nostoc*-*Gunnera* symbiosis. New Phytol 122:379–400.

46. Mutalipassi M, Riccio G, Mazzella V, Galasso C, Somma E, Chiarore A, de Pascale D, Zupo V. 2021. Symbioses of Cyanobacteria in Marine Environments: Ecological Insights and Biotechnological Perspectives. Mar Drugs 19:227.

47. Val-Calvo J, Vázquez-Boland JA. 2023. *Mycobacteriales* taxonomy using network analysis-aided, context-uniform phylogenomic approach for non-subjective genus demarcation. mBio 14:e02207–23.

48. Morgado S, Ramos N de V, Pereira BB do N, Freitas F, Fonseca ÉL da, Vicente AC. 2022. Multidrug-resistant *Mycolicibacterium fortuitum* infection in a companion cat (*Felis silvestris catus*) in Brazil. Access Microbiology 4:000317.

49. Antunes LC, Poppleton D, Klingl A, Criscuolo A, Dupuy B, Brochier-Armanet C, Beloin C, Gribaldo S. 2016. Phylogenomic analysis supports the ancestral presence of LPS-outer membranes in the Firmicutes. eLife 5:e14589.

50. Brown RF. 2016. Investigating the Evolutionary Origin and Cell Biology of Negativicutes. University of Warwick, Warwick.

51. Rahimlou S, Bahram M, Tedersoo L. 2021. Phylogenomics reveals the evolution of root nodulating alpha- and beta-Proteobacteria (rhizobia). Microbiological Research 250:126788.

52. Lavin M, Herendeen PS, Wojciechowski MF. 2005. Evolutionary Rates Analysis of *Leguminosae* Implicates a Rapid Diversification of Lineages during the Tertiary. Systematic Biology 54:575–594.

53. Travin DY, Jouan R, Vigouroux A, Inaba-Inoue S, Lachat J, Haq F, Timchenko T, Sutormin D, Dubiley S, Beis K, Moréra S, Severinov K, Mergaert P. 2023. Dual-Uptake Mode of the Antibiotic Phazolicin Prevents Resistance Acquisition by Gram-Negative Bacteria. mBio 14:e00217–23.

54. Marlow VL, Haag AF, Kobayashi H, Fletcher V, Scocchi M, Walker GC, Ferguson GP. 2009. Essential role for the BacA protein in the uptake of a truncated eukaryotic peptide in *Sinorhizobium meliloti*. J Bacteriol 191:1519–1527.

55. Wells DH, Long SR. 2002. The *Sinorhizobium meliloti* stringent response affects multiple aspects of symbiosis. Molecular microbiology 43:1115–1127.

56. Langmead B, Salzberg SL. 2012. Fast gapped-read alignment with Bowtie 2. Nature Methods 2012 9:4 9:357–359.

57. Robinson JT, Thorvaldsdóttir H, Winckler W, Guttman M, Lander ES, Getz G, Mesirov JP. 2011. Integrative genomics viewer. Nature Biotechnology 2011 29:1 29:24–26.

58. Finan TM, Kunkel B, De Vos GF, Signer ER. 1986. Second symbiotic megaplasmid in *Rhizobium meliloti* carrying exopolysaccharide and thiamine synthesis genes. J Bacteriol 167:66.

59. Jensen HL, Jensen HL. 1942. Nitrogen fixation in leguminous plants. I. General characters of root-nodule bacteria isolated from species of *Medicago* and *Trifolium* in Australia. Proceedings of the Linnean Society of New South Wales 67:98–108.

60. diCenzo GC, Mengoni A, Perrin E. 2019. Chromids Aid Genome Expansion and Functional Diversification in the Family *Burkholderiaceae*. Molecular Biology and Evolution 36:562–574.

61. Johnson LS, Eddy SR, Portugaly E. 2010. Hidden Markov model speed heuristic and iterative HMM search procedure. BMC Bioinformatics 11:431.

62. Sievers F, Wilm A, Dineen D, Gibson TJ, Karplus K, Li W, Lopez R, McWilliam H, Remmert M, Söding J, Thompson JD, Higgins DG. 2011. Fast, scalable generation of high-quality protein multiple sequence alignments using Clustal Omega. Mol Syst Biol 7:539.

63. Katoh K, Standley DM. 2013. MAFFT Multiple Sequence Alignment Software Version 7: Improvements in Performance and Usability. Molecular Biology and Evolution 30:772–780.

64. Notredame C, Higgins DG, Heringa J. 2000. T-Coffee: A novel method for fast and accurate multiple sequence alignment. J Mol Biol 302:205–217.

65. Capella-Gutiérrez S, Silla-Martínez JM, Gabaldón T. 2009. trimAl: a tool for automated alignment trimming in large-scale phylogenetic analyses. Bioinformatics 25:1972–1973.

66. Minh BQ, Schmidt HA, Chernomor O, Schrempf D, Woodhams MD, von Haeseler A, Lanfear R. 2020. IQ-TREE 2: New Models and Efficient Methods for Phylogenetic Inference in the Genomic Era. Molecular Biology and Evolution 37:1530–1534.

67. Kalyaanamoorthy S, Minh BQ, Wong TKF, von Haeseler A, Jermiin LS. 2017. ModelFinder: fast model selection for accurate phylogenetic estimates. 6. Nat Methods 14:587–589.

68. Guindon S, Dufayard J-F, Lefort V, Anisimova M, Hordijk W, Gascuel O. 2010. New algorithms and methods to estimate maximum-likelihood phylogenies: assessing the performance of PhyML 3.0. Syst Biol 59:307–321.

69. Letunic I, Bork P. 2021. Interactive Tree Of Life (iTOL) v5: an online tool for phylogenetic tree display and annotation. Nucleic Acids Research 49:W293–W296.

70. Oberg N, Zallot R, Gerlt JA. 2023. EFI-EST, EFI-GNT, and EFI-CGFP: Enzyme Function Initiative (EFI) Web Resource for Genomic Enzymology Tools. Journal of Molecular Biology 435:168018.

71. Zallot R, Oberg N, Gerlt JA. 2019. The EFI Web Resource for Genomic Enzymology Tools: Leveraging Protein, Genome, and Metagenome Databases to Discover Novel Enzymes and Metabolic Pathways. Biochemistry 58:4169–4182.

72. Shannon P, Markiel A, Ozier O, Baliga NS, Wang JT, Ramage D, Amin N, Schwikowski B, Ideker T. 2003. Cytoscape: a software environment for integrated models of biomolecular interaction networks. Genome Res 13:2498–2504.

73. Wu M, Scott AJ. 2012. Phylogenomic analysis of bacterial and archaeal sequences with AMPHORA2. Bioinformatics 28:1033–1034.

