## Supplementary material for "Taxonomic distribution of SbmA/BacA and BacA-like antimicrobial peptide transporters suggests independent recruitment and convergent evolution in host-microbe interactions": File_S1

**This PDF file includes:**

Tables S1 to S2 (pages 2-4)
Figures S1 to S4 (pages 5-8)
Legends for Datasets S1 to S2 (page 9)

**Other supplementary materials for this manuscript include the following:**

Datasets S1 to S2

**Table S1.** Shoot dry weights of legumes inoculated with a *Sinorhizobium meliloti* Δ*bacA* mutant and *S. meliloti* Δ*bacA* mutants expressing various *sbmA*-like proteins *in trans*.

| ***S. meliloti* genotype *** | **Mean shoot dry weight ± standard deviation (mg/plant)** |
|---|---|
| ***Medicago sativa* (alfalfa)** | |
| Wildtype Rm1021 | 66.7 ± 7.1 |
| Δ*bacA* empty vector control | 7.6 ± 1.1 |
| *Sinorhizobium meliloti* 1021 *bacA* | 68.6 ± 4.9 |
| *Polymorphum gilvum bacA* | 8.8 ± 1.7 |
| *Bradyrhizobium* sp. ORS285 *bclA* | 7.1 ± 0.6 |
| *Succinivibrio dextrinosolvens bclA* | 7.7 ± 1.1 |
| *Basilea psittacipulmonis bclA* | 10.7 ± 1.2 |
| *Polaromonas naphthalenivorans bclA* | 8.8 ± 0.8 |
| *Synechococcus elongatus bclA* | 6.1 ± 0.9 |
| *Methylomusa anaerophila bclA* | 8 ± 1.4 |
| *Cyanobacterium aponinum bclA* | 7.1 ± 0.2 |
| *Eikenella exigua bclA*-like | 6.7 ± 1.3 |
| *Phyllobacterium zundukense exsE* | 7.7 ± 1.2 |
| Uninoculated control | 6.5 ± 1.5 |
| ***Melilotus officinalis* (yellow-blossom sweet clover)** | |
| Wild type Sm1021 | 72.2 ± 4.5 |
| Δ*bacA* | 6.8 ± 1.3 |
| *Sinorhizobium meliloti* 1021 *bacA* | 35.6 ± 8.7 |
| *Polymorphum gilvum bacA* | 5.0 ± 0.6 |
| *Bradyrhizobium* sp. ORS285 *bclA* | 3.1 ± 0.7 |
| *Succinivibrio dextrinosolvens bclA* | 6.3 ± 1.3 |
| *Basilea psittacipulmonis bclA* | 5.2 ± 0.2 |
| *Polaromonas naphthalenivorans bclA* | 3.5 ± 1.9 |
| *Synechococcus elongatus bclA* | 6.8 ± 1.9 |
| *Methylomusa anaerophila bclA* | 6.9 ± 0.6 |
| *Cyanobacterium aponinum bclA* | 4.6 ± 0.2 |
| *Eikenella exigua bclA*-like | 7.2 ± 0.4 |
| *Phyllobacterium zundukense exsE* | 3.6 ± 0.2 |
| Uninoculated | 6 ± 1.1 |

* Strains included the wildtype *S. meliloti* strain Rm1021, a Rm1021 Δ*bacA* derivative, and Δ*bacA* derivatives expressing the genes from the indicated organisms *in trans*.

**Table S2.** Bacterial strains and plasmids

| Strain/Plasmid | Characteristics | Reference |
|---|---|---|
| ***Sinorhizobium meliloti*** | | |
| Sm1021 | Wild type SU47 *str-21*; $Sm^R$ | Meade *et al*. (1982) |
| SmGF1 | Sm1021 *bacA*654::Spc (Δ*bacA* null); $Sm^R$ $Sp^R$ | Ferguson *et al*. (2002) |
| Sm1021Δ*bacA*Ω*yejA* | Sm1021 Δ*bacA* Ω*yejA*; $Sm^R$ $Sp^R$ $Km^R$ | Travin DY *et al*. (2023) |
| SmGQ0019 | SmGF1 (pGQ0037); $Sm^R$ $Sp^R$ $Tc^R$ | This study |
| SmGQ0020 | SmGF1 (pGQ0038); $Sm^R$ $Sp^R$ $Tc^R$ | This study |
| SmGQ0021 | SmGF1 (pGQ0039); $Sm^R$ $Sp^R$ $Tc^R$ | This study |
| SmGQ0022 | SmGF1 (pGQ0040); $Sm^R$ $Sp^R$ $Tc^R$ | This study |
| SmGQ0023 | SmGF1 (pGQ0041); $Sm^R$ $Sp^R$ $Tc^R$ | This study |
| SmGQ0024 | SmGF1 (pGQ0042); $Sm^R$ $Sp^R$ $Tc^R$ | This study |
| SmGQ0025 | SmGF1 (pGQ0043); $Sm^R$ $Sp^R$ $Tc^R$ | This study |
| SmGQ0026 | SmGF1 (pGQ0044); $Sm^R$ $Sp^R$ $Tc^R$ | This study |
| SmGQ0027 | SmGF1 (pGQ0045); $Sm^R$ $Sp^R$ $Tc^R$ | This study |
| SmGQ0028 | SmGF1 (pGQ0046); $Sm^R$ $Sp^R$ $Tc^R$ | This study |
| SmGQ0029 | SmGF1 (pRF771:*bclA*-ORS285); $Sm^R$ $Sp^R$ $Tc^R$ | This study |
| SmGQ0030 | SmGF1 (pRF771); $Sm^R$ $Sp^R$ $Tc^R$ | This study |
| SmGQ0031 | SmGF1 (pRF771::*bacA*-Sm1021); $Sm^R$ $Sp^R$ $Tc^R$ | This study |
| SmGQ0032 | SmGF1 (pRF771::*bclA*-USDA110); $Sm^R$ $Sp^R$ $Tc^R$ | This study |
| SmYjaGQ0019 | Sm1021Δ*bacA*Ω*yejA* (pGQ0037); $Sm^R$ $Tc^R$ $Sp^R$ $Km^R$ | This study |
| SmYjaGQ0020 | Sm1021Δ*bacA*Ω*yejA* (pGQ0038); $Sm^R$ $Tc^R$ $Sp^R$ $Km^R$ | This study |
| SmYjaGQ0021 | Sm1021Δ*bacA*Ω*yejA* (pGQ0039); $Sm^R$ $Tc^R$ $Sp^R$ $Km^R$ | This study |
| SmYjaGQ0022 | Sm1021Δ*bacA*Ω*yejA* (pGQ0040); $Sm^R$ $Tc^R$ $Sp^R$ $Km^R$ | This study |
| SmYjaGQ0023 | Sm1021Δ*bacA*Ω*yejA* (pGQ0041); $Sm^R$ $Tc^R$ $Sp^R$ $Km^R$ | This study |
| SmYjaGQ0024 | Sm1021Δ*bacA*Ω*yejA* (pGQ0042); $Sm^R$ $Tc^R$ $Sp^R$ $Km^R$ | This study |
| SmYjaGQ0025 | Sm1021Δ*bacA*Ω*yejA* (pGQ0043); $Sm^R$ $Tc^R$ $Sp^R$ $Km^R$ | This study |
| SmYjaGQ0026 | Sm1021Δ*bacA*Ω*yejA* (pGQ0044); $Sm^R$ $Tc^R$ $Sp^R$ $Km^R$ | This study |
| SmYjaGQ0027 | Sm1021Δ*bacA*Ω*yejA* (pGQ0045); $Sm^R$ $Tc^R$ $Sp^R$ $Km^R$ | This study |
| SmYjaGQ0028 | Sm1021Δ*bacA*Ω*yejA* (pGQ0046); $Sm^R$ $Tc^R$ $Sp^R$ $Km^R$ | This study |
| SmYjaGQ0029 | Sm1021Δ*bacA*Ω*yejA* (pRF771::*bclA*-ORS285); $Sm^R$ $Tc^R$ $Sp^R$ $Km^R$ | This study |
| SmYjaGQ0030 | Sm1021Δ*bacA*Ω*yejA* (pRF771); $Sm^R$ $Tc^R$ $Sp^R$ $Km^R$ | This study |
| SmYjaGQ0031 | Sm1021Δ*bacA*Ω*yejA* (pRF771::*bacA*-Sm1021); $Sm^R$ $Tc^R$ $Sp^R$ $Km^R$ | This study |
| SmYjaGQ0032 | Sm1021Δ*bacA*Ω*yejA* (pRF771::*bacA*-USDA110); $Sm^R$ $Tc^R$ $Sp^R$ $Km^R$ | This study |
| ***Escherichia coli*** | | |
| MT616 | MM294A *recA-56* (pRK600), mobilizer; $Cm^R$ | Finan *et al*. (1986) |
| HB101 | F–, *thi*-1, *hsdS*20 ($r_B^-$, $m_B^-$), *supE*44, *recA*13, *ara*-14, *leuB*6, *proA*2, *lacY*1, *galK*2, *rpsL*20, *xyl*-5, *mtl*-1; $Sm^R$ | Boyer & Roulland-Dussoix (1969) |
| **Plasmids** | | |
| pRF771 | pTE3 derivative with *trp* promoter; $Tc^R$ | Wells and Long (2002) |
| pRF771::*bacA*-Sm1021 | pRF771::*Sinorhizobium meliloti* 1021 *bacA*; $Tc^R$ | Haag *et al*. (2011) |
| pRF771:*bclA*-ORS285 | pRF771::*Bradyrhizobium* sp. ORS285 *bclA*; $Tc^R$ | Guefrachi *et al*. (2015) |
| pRF771::*bclA*-USDA110 | pRF771::*Bradyrhizobium diazoefficiens* USDA110 *bclA*, $Tc^R$ | Barrière *et al*. (2017) |
| pGQ0037 | pRF771::*Succinivibrio dextrinosolvens bclA*; $Tc^R$ | This study |
| pGQ0038 | pRF771::*Polymorphum gilvum bacA*; $Tc^R$ | This study |
| pGQ0039 | pRF771::*Basilea psittacipulmonis bclA*; $Tc^R$ | This study |
| pGQ0041 | pRF771::*Polaromonas naphthalenivorans bclA*; $Tc^R$ | This study |
| pGQ0042 | pRF771::*Phyllobacterium zundukense exsE*; $Tc^R$ | This study |
| pGQ0043 | pRF771::*Eikenella exigua bclA*-like gene; $Tc^R$ | This study |
| pGQ0044 | pRF771::*Synechococcus elongatus bclA*; $Tc^R$ | This study |
| pGQ0045 | pRF771::*Methylomusa anaerophila bclA*; $Tc^R$ | This study |
| pGQ0046 | pRF771::*Cyanobacterium aponinum bclA*; $Tc^R$ | This study |

Sm – streptomycin; Sp – spectinomycin; Km – kanamycin; Tc – tetracycline; Cm – chloramphenicol


Barrière, Q., Guefrachi, I., Gully, D., Lamouche, F., Pierre, O., Fardoux, J., Chaintreuil, C., Alunni, B., Timchenko, T., Giraud, E., and Mergaert, P. 2017. Integrated roles of BclA and DD-carboxypeptidase 1 in *Bradyrhizobium* differentiation within NCR-producing and NCR-lacking root nodules. Sci Rep **7**(1): 9063. doi:10.1038/s41598-017-08830-0.

Boyer, H.W., and Roulland-dussoix, D. 1969. A complementation analysis of the restriction and modification of DNA in *Escherichia coli*. J Mol Biol **41**(3): 459–472. doi:10.1016/0022-

2836(69)90288-5.

Ferguson, G.P., Roop, R.M., and Walker, G.C. 2002. Deficiency of a *Sinorhizobium meliloti bacA* mutant in alfalfa symbiosis correlates with alteration of the cell envelope. J Bacteriol **184**(20): 5625–5632. doi:10.1128/JB.184.20.5625-5632.2002.

Finan, T.M., Kunkel, B., De Vos, G.F., and Signer, E.R. 1986. Second symbiotic megaplasmid in *Rhizobium meliloti* carrying exopolysaccharide and thiamine synthesis genes. J Bacteriol **167**(1): 66–72.

Guefrachi, I., Pierre, O., Timchenko, T., Alunni, B., Barrière, Q., Czernic, P., Villaécija-Aguilar, J.-A., Verly, C., Bourge, M., Fardoux, J., Mars, M., Kondorosi, E., Giraud, E., and Mergaert, P. 2015. *Bradyrhizobium* BclA is a peptide transporter required for bacterial differentiation in symbiosis with *Aeschynomene* legumes. Mol Plant Microbe Interact **28**(11): 1155–1166. doi:10.1094/MPMI-04-15-0094-R.

Haag, A.F., Baloban, M., Sani, M., Kerscher, B., Pierre, O., Farkas, A., Longhi, R., Boncompagni, E., Hérouart, D., Dall'Angelo, S., Kondorosi, E., Zanda, M., Mergaert, P., and Ferguson, G.P. 2011. Protection of *Sinorhizobium against* host cysteine-rich antimicrobial peptides is critical for symbiosis. PLOS Biol **9**(10): e1001169. doi:10.1371/journal.pbio.1001169.

Meade, H.M., Long, S.R., Ruvkun, G.B., Brown, S.E., and Ausubel, F.M. 1982. Physical and genetic characterization of symbiotic and auxotrophic mutants of *Rhizobium meliloti* induced by transposon Tn*5* mutagenesis. J Bacteriol **149**(1): 114–122.

Travin, D.Y., Jouan, R., Vigouroux, A., Inaba-Inoue, S., Lachat, J., Haq, F., Timchenko, T., Sutormin, D., Dubiley, S., Beis, K., Moréra, S., Severinov, K., and Mergaert, P. 2023. Dual-uptake mode of the antibiotic phazolicin prevents resistance acquisition by Gram-negative bacteria. mBio **14**(2): e0021723. doi:10.1128/mbio.00217-23.

Wells, D.H., and Long, S.R. 2002. The *Sinorhizobium meliloti* stringent response affects multiple aspects of symbiosis. Mol Microbiol **43**(5): 1115–1127. doi:10.1046/j.1365-2958.2002.02826.x.

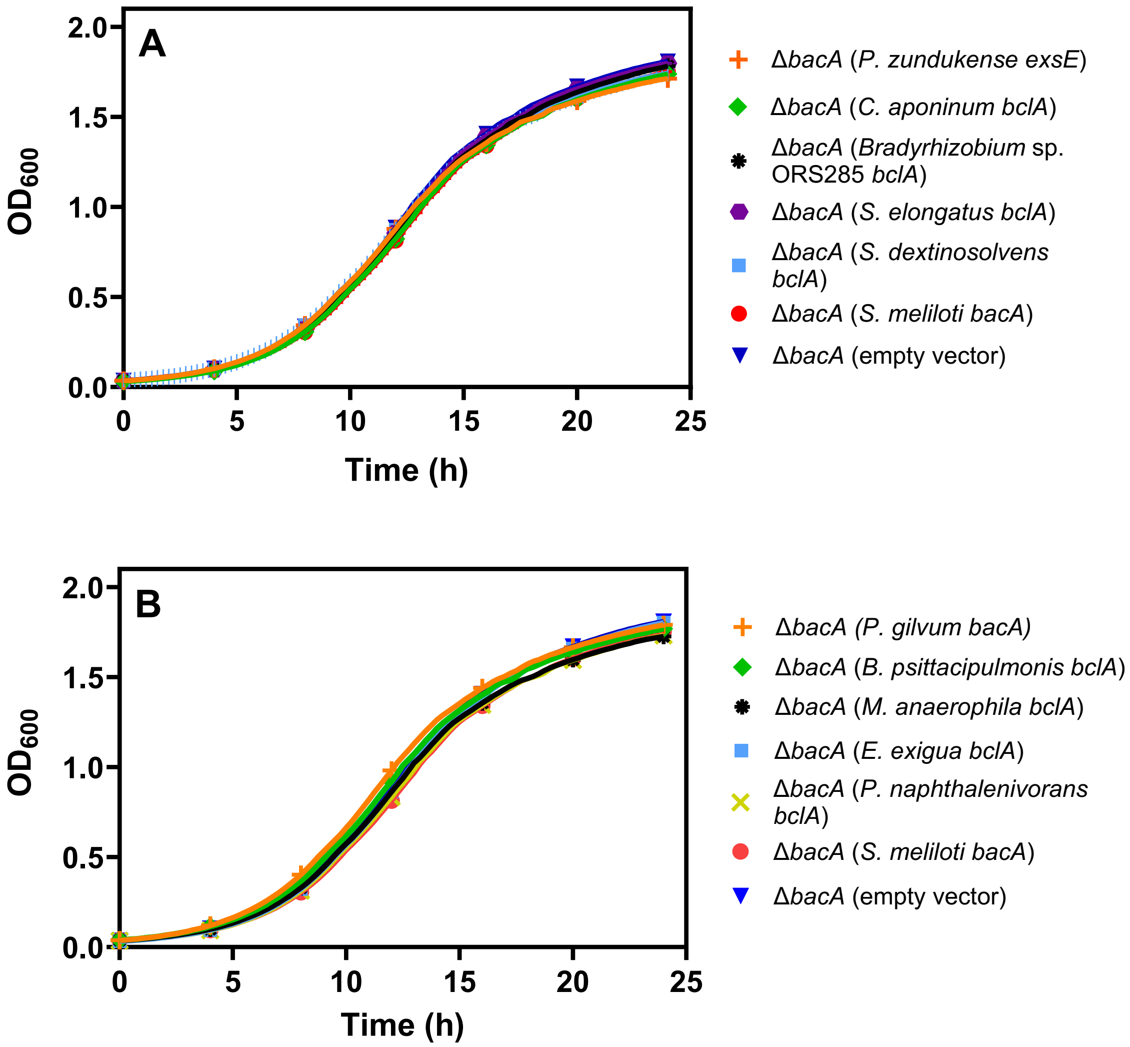


**Figure S1. Growth of *Sinorhizobium meliloti* strains in LBmc.** The growth of various *S. meliloti* strains, as measured by OD600, in LBmc is shown over a 24-hour period. Each point represents the mean of triplicate wells, with error bars depicting standard deviation. The Δ*bacA* strain represents the *S. meliloti* Δ*bacA* mutant carrying an empty vector, while all other strains are named according to the species of origin of the gene expressed *in trans* in the *S. meliloti* Δ*bacA* background. The experiment was replicated three independent times, and data from a representative experiment is shown.

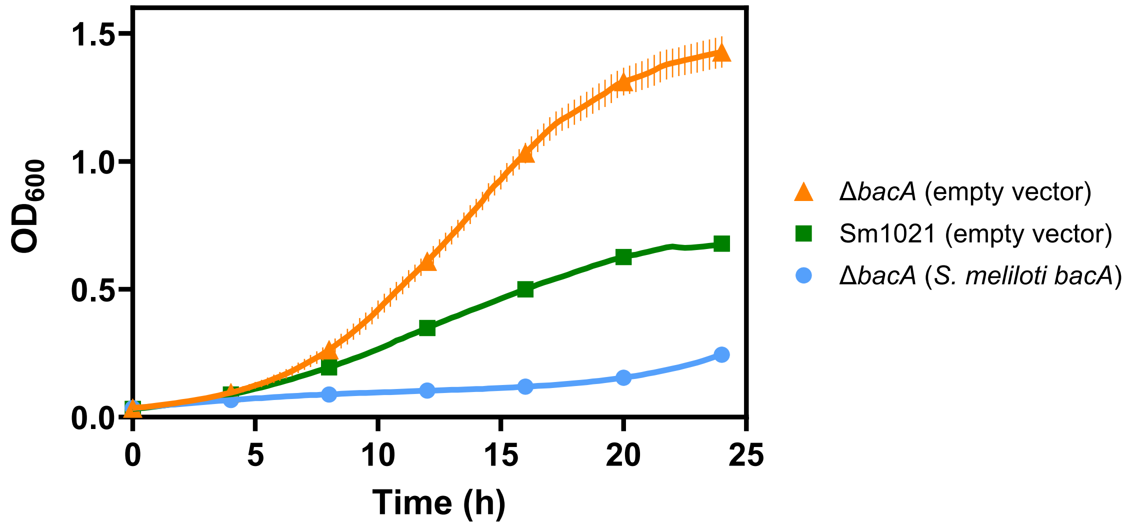


**Figure S2. Effect of expressing *bacA in trans* on the gentamicin sensitivity of *Sinorhizobium meliloti*.** The growth of three *S. meliloti* strains, as measured by OD600, in the presence of 20 µg/mL of gentamicin is shown over a 24-hour period. Growth profiles are shown for wildtype *meliloti* Sm1021 harbouring an empty expression vector (blue), and a *S. meliloti* Δ*bacA* mutant expressing (blue) or not (orange) the *S. meliloti bacA* gene *in trans*. Each point represents the mean of triplicate wells, with error bars depicting standard deviation. The experiment was replicated three independent times, and data from a representative experiment is shown.

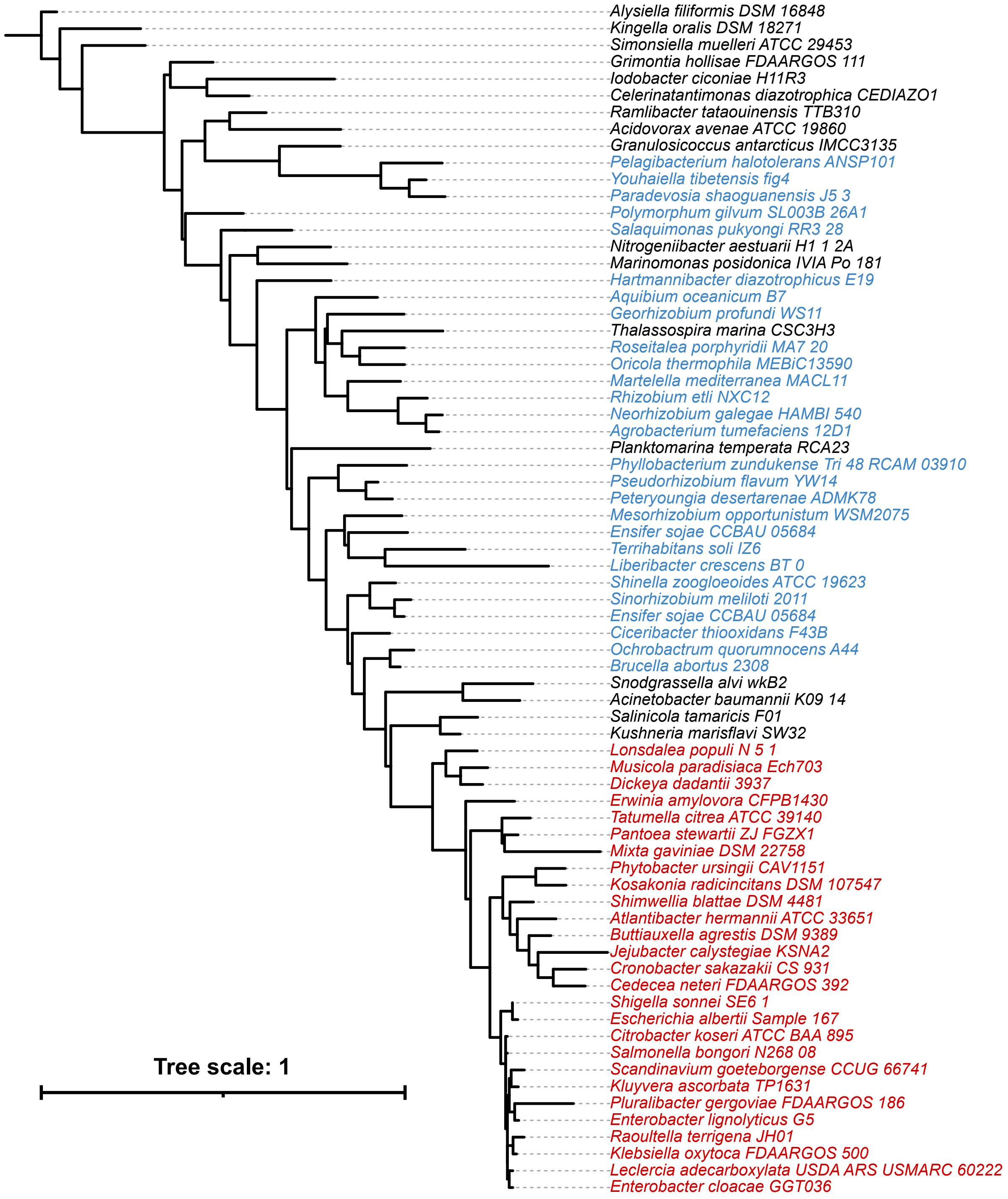


**Figure S3. Maximum-likelihood phylogeny of SbmA/BacA proteins.** A subtree of the maximum likelihood phylogeny of SbmA/BacA-like proteins of Figure 1 is shown. This subtree is limited to the 71 proteins classified as BacA. Protein encoded by species of the order *Enterobacterales* are shown in red, while proteins of the order *Hyphomicrobiales* are shown in blue. The scale bar represents the average number of amino acid substitutions per site. An interactive version of this phylogeny, with node support values, is provided through iTol (https://itol.embl.de/shared/1IAjjFrHYGLI9).

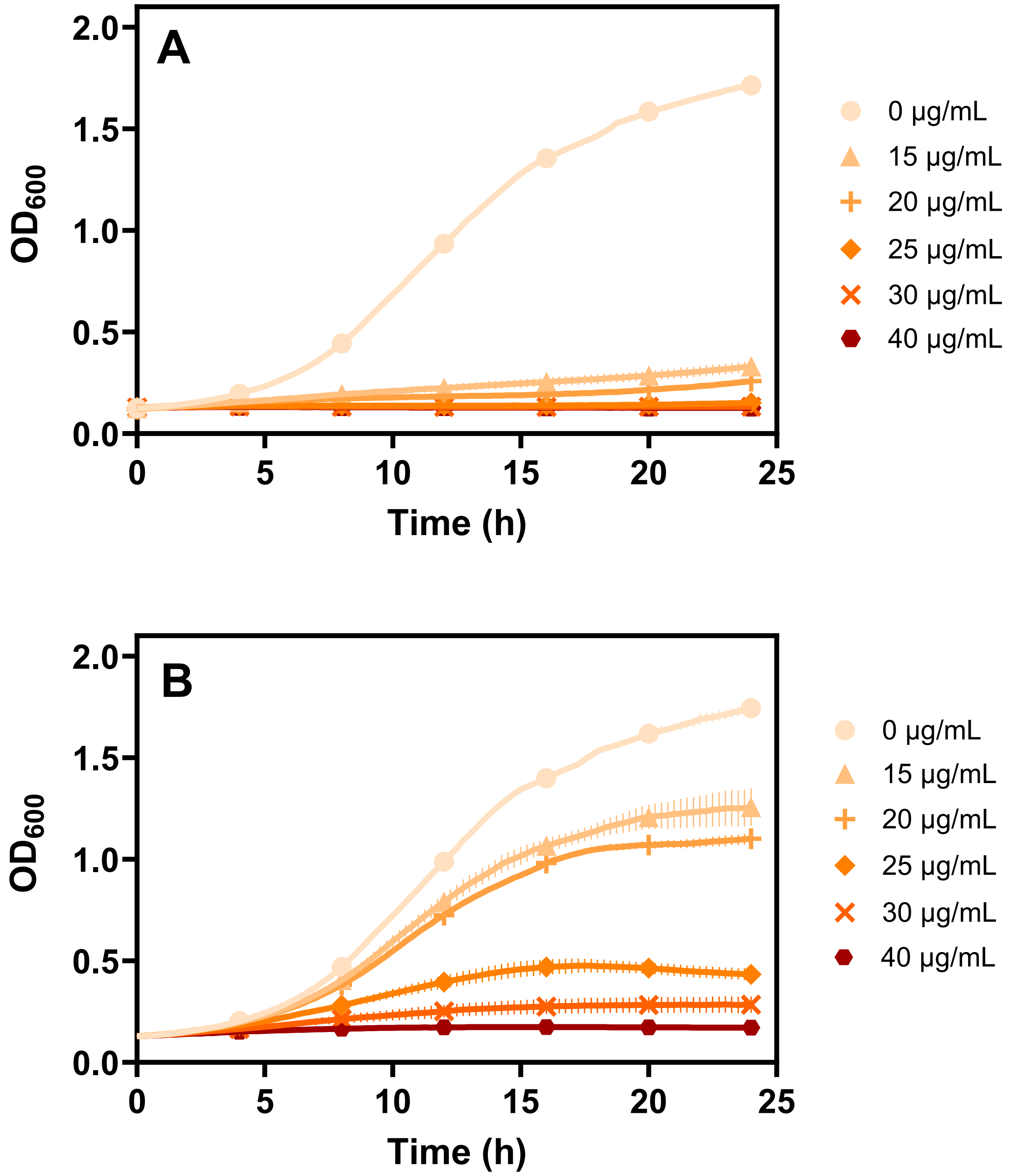


**Figure S4. Effect of gentamicin on the growth of *Sinorhizobium meliloti*.** The growth of *S. meliloti* in LBmc containing various concentrations of gentamicin, as indicated in the legend. Growth profiles are shown for (**A**) *S. meliloti* Δ*bacA* carrying an empty expression vector, and (**B**) *S. meliloti* Δ*bacA* expressing the *S. meliloti bacA* gene *in trans*. Each point represents the mean of triplicate wells, with error bars depicting standard deviation. The experiment was replicated three independent times, and data from a representative experiment is shown.

### DATASET LEGENDS

**Dataset S1.** Metadata for the 1,255 bacterial genomes used in this study, including ftp links to download the genomes from the National Center for Biotechnology Information (NCBI) database. **(.TSV file; tab-delimited)**

**Dataset S2.** The sequences, locus tag, and organism or origin for the 366 proteins included in the phylogeny of Figure 1. The first column indicates the nine genes that were codon optimized for *Sinorhizobium meliloti* and synthesized. **(.TSV file; tab-delimited)**
